# Dehydration-Responsive Left-Handed G-Quadruplexes Expand the Sequence Repertoire

**DOI:** 10.64898/2026.10.05.756892

**Authors:** Yun Chen, Andrew Hendrickson, Julien Gros, Eric Xing, Eleanor R. Windle, Jiawei Wang, Robert M. Edkins, Susan J. Quinn, Liliya A. Yatsunyk, Jean-Louis Mergny

## Abstract

G-quadruplexes (G4s) are widely studied noncanonical nucleic-acid structures that carry important biological functions. Typically, G4s form from G-rich sequences containing at least four tracts of more than two consecutive guanines, yet a plethora of unconventional G4-forming sequences have been identified, including sequences that fold into G4s only under specific conditions. These G4 states remain poorly resolved. Here we identify *dehydration-responsive left-handed G4s* (drLHG4s), a class of left-handed G4 (LHG4) structures accessible under dehydrating conditions. Using d(GT)_n_ repeats, we show that left-handed folding depends on repeat length, salt condition and solvent conditions. Systematic mutational analysis reveals a broader but register-constrained sequence space, while high-resolution crystal structures of d-GTGGTGTGTGGTG, a sequence that folded robustly under dehydrating conditions, establish a bulged LHG4 architecture and reveal a distinctive hydration sphere. We further show that selected peptides and a cationic zinc phthalocyanine derivative, ZnTEG^Q^PyrPc, containing four triethylene glycol methyl ether tails can induce LHG4 states. Together, our findings reveal a dehydration-responsive G4 landscape and expand the experimentally accessible sequence space of LHG4 folding.

## Introduction

G-quadruplexes (G4s) are noncanonical structures formed by guanine-rich DNA or RNA. They have been implicated in the regulation of transcription and replication and in genome instability.^1-7^ Their diversity is well established. Parallel, antiparallel and hybrid topologies are widely documented,^6–9^ and additional variation arises from tetrad number,^10–14^ bulges,^15^ non-G tetrads,^16–20^ capping base pairs or triads,^21–23^ and junctional organization.^24–27^ These features alter exposed surfaces and molecular recognition.^28,29^ However, this diversity has not produced a clear biological hierarchy. In most studies, G-rich sequences are classified primarily as G4-forming or non-G4-forming.^30^ Similarly, cellular imaging and genome-wide mapping approaches usually report G4 formation rather than topology.^31–33^ Such classifications may overlook sequences whose G4 folding becomes accessible only under particular environmental conditions. Namely, structurally distinct G4 motifs often remain grouped together at the biological level, even if individual sequences can switch between folds depending on ions,^20,34–36^ hydration,^37–40^ cosolutes,^39–41^ and ligands.^42,43^ Cellular microenvironments can alter G4 stability and its dependence on solution conditions, motivating the exploration of G4 folding beyond standard dilute buffers.^44^

Double-stranded nucleic acids provide a useful contrast. Z-DNA for instance is defined not only by its left-handed structure but also by its activation requirements. These include sequence context,^45^ negative supercoiling,^46,47^ salt conditions,^48,49^ chemical modification,^50,51^ and selective protein recognition.^52,53^ We therefore asked whether analogous condition-dependent access might exist for specific G4 folds.

Left-handed G-quadruplexes (LHG4s) provide a natural entry point. They are distinct G4 architectures characterized by reversed backbone progression, characteristic tetrad stacking and defined loop organization, rather than variants of conventional right-handed G4 (RHG4) topologies.^54–58^ Structural studies have established LHG4s as a genuine fold class and identified two minimal sequence frameworks for LHG4 formation: d-GTGGTGGTGGTG (*Motif1*) and d-GGTGGTGGTGTG (*Motif2*).^54,55,57^ These studies also suggest that left-handed folding is a constrained, motif-dependent G4 folding mode.

Here we revisit this boundary using d(GT)_n_ repeats and their variants (Table S1). We show that left-handed folding extends beyond known motifs, and that the expanded landscape is not uniform. We refer to this broader condition-accessible branch as dehydration-responsive left-handed G-quadruplexes (drLHG4s). Our findings expand the known conformational landscape of G4 folding by revealing a left-handed-related branch that is poorly sampled under standard aqueous conditions. Using spectroscopy, crystallography, sequence analysis, and molecular induction, we establish an experimental framework for probing this hidden layer of G4 structural plasticity.

## Results

### d(GT)_n_ repeats access LHG4 folding through a dehydration-responsive switch

Structurally characterized minimal LHG4-forming sequences are built around two closely related motifs, d-GTGGTGGTGGTG (*Motif1*) and d-GGTGGTGGTGTG (*Motif2*).^54,55^ Both feature parallel strand orientation, left-handed backbone progression, thymine caps, single-nucleotide edge loops and higher-order assembly of two two-tetrad LHG4 blocks, together defining the LHG4 folding mode.^57,59^

d(GT)_n_ and d(GA)_n_ repeats are abundant in genomes,^60–63^ and increasingly recognized as biologically active elements.^63–65^ Short tandem repeats are enriched in regulatory regions and can tune transcription factor binding and gene expression.^66^ They have also been proposed as potential sources of noncanonical quadruplex folds.^67^ Their sequence similarity to known LHG4 motifs prompted us to test whether they also form LHG4s.

Differences in water organization between Z-DNA and B-DNA and between LHG4 and RHG4 prompted us to ask whether reduced water activity could promote LHG4 folding.^68–69^ We first tested d(GT)_n_ repeats in the presence of PEG200, which contains on average four ethylene glycol units. Low-molecular-weight PEGs such as PEG200 are commonly used to lower water activity, whereas higher-molecular-weight PEGs more strongly contribute to excluded-volume effects.^39,40,70–72^

To test the effect of dehydrating conditions on LHG4 formation, we first used PEG200 as a low-molecular-weight cosolvent. In the absence of PEG200, a representative sequence d(GT)_12_ showed no clear G4-like CD signature, whereas PEG200 titration from 0 to 75% (v/v) induced a LHG4 CD profile, characterized by the growth of a positive band near 245 nm and a negative band near 265 nm, matching the spectral signature reported for structurally characterized LHG4s (Figure 1A).^54–56^ Truncation analysis starting from d(GT)_16_ down to d(GT)_5_ showed that d(GT)_7.5_ was the shortest repeat exhibiting this behavior (Figure 1B, 1C). This sequence contains eight guanines, the minimal number required for the formation of two complete G-quartets. The same ability to form LHG4 structures induced by the presence of PEG200 and the same length requirement was observed in the d(GA)_n_ series (n between 6 and 12; Table S1, Figure S1, S2).

**Figure 1.**
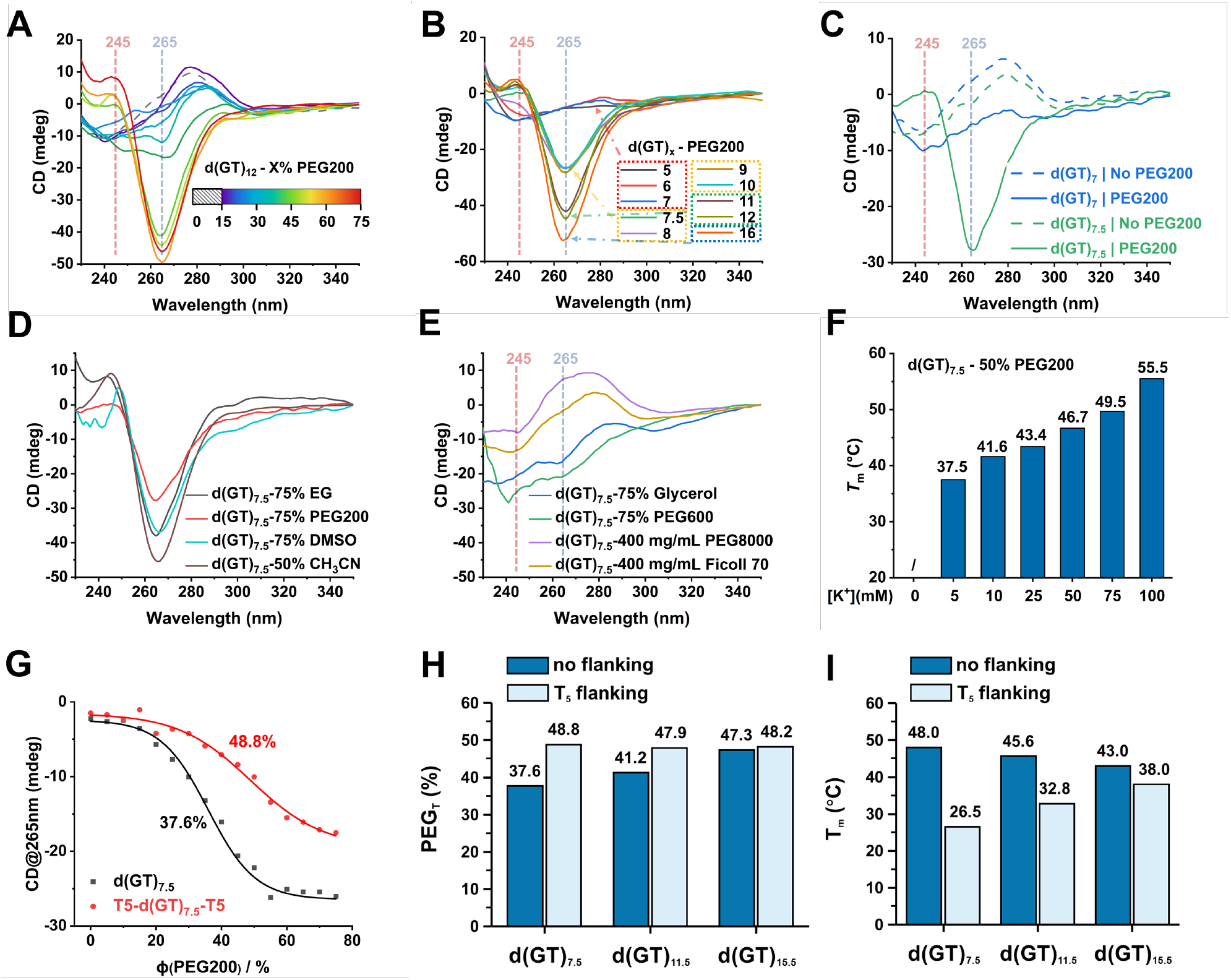
Dehydration-responsive left-handed folding in d(GT)_n_ repeats. Unless otherwise stated, all CD spectra were recorded at 5 μM DNA strand concentration. PEG200 and other liquid cosolvents are reported as % v/v. (**A**) CD spectra of d(GT)_12_ measured at increasing PEG200 concentrations, showing progressive induction of the characteristic LHG4 signature. Dashed vertical lines indicate characteristic wavelengths (245 and 265 nm). (**B**) CD spectra of d(GT)_n_ repeats of different lengths measured in 50% PEG200. (**C**) CD spectra of d(GT)_7_ and d(GT)_7.5_ recorded in the absence or presence of 75% PEG200, shown as dashed and solid lines, respectively. (**D**) CD spectra of d(GT)_7.5_ measured in 75% ethylene glycol, PEG200, DMSO, or 50% acetonitrile. (**E**) CD spectra of d(GT)_7.5_ measured in 75% glycerol, 75% PEG600, 400 mg/mL PEG8000, or 400 mg/mL Ficoll 70. (**F**) Melting temperatures of d(GT)_7.5_ in 50% PEG200 at increasing K^+^ concentrations. (**G**) Determination of PEG_T_ from PEG200 titrations monitored by CD at 265 nm and 25 °C for d(GT)_7.5_ and T_5_-d(GT)_7.5_-T_5_. PEG_T_ values are indicated at the midpoints of the folding transitions. (**H-I**) T_5_ flanking sequences disfavor LHG4 folding: (**H**) Comparison of PEG_T_ values for d(GT)_7.5_, d(GT)_11.5_ and d(GT)_15.5_, with or without T_5_ flanking sequences. (**I**) Comparison of *T*_m_ values for d(GT)_7.5_, d(GT)_11.5_ and d(GT)_15.5_, with or without T_5_ flanking sequences.

We next compared PEG200 with other cosolvents and crowding agents to distinguish dehydration-related effects from excluded-volume crowding. Ethylene glycol, DMSO and acetonitrile induced LHG4-like folding (Figure 1D). In agreement with our CD results, ^1^H NMR spectra of d(GT)_7.5_ in 50% DMSO (v/v) showed imino resonances consistent with Hoogsteen hydrogen bonding (Figure S3). On the contrary, higher-molecular-weight PEG600 and PEG8000, glycerol, and Ficoll 70 did not induce LHG4 folding (Figure 1D, E).^73^ This contrast argues against a simple excluded-volume mechanism (“crowding effect”) and suggests that productive folding requires solvent conditions capable of remodeling local hydration around nascent tetrad-loop interfaces. PEG200 promoted folding at higher water activity than DMSO, whereas glycerol failed to induce comparable folding despite reaching a similar water-activity range (Figure S4, S5), indicating that bulk water activity alone does not determine LHG4 formation. Solvent-specific interactions, potentially involving local hydration, may also contribute.

LHG4 folding also showed a strong preference for K^+^. Increasing K^+^ concentration progressively stabilized the structure of d(GT)_7.5_ (Figure 1F, S6). Na^+^ was much less effective, and Mg^2+^ or Ca^2+^ did not support LHG4 formation in the 0-100 mM cation concentration range (Figure S6). This K^+^ preference is consistent with canonical G-quartet channel stabilization, in which potassium ions are coordinated by inward-facing guanine O6 carbonyls between stacked tetrads. Ion size, coordination geometry and dehydration energetics all contribute to this preference.^74,75^

We quantified the dehydration requirement using PEG_T_, an operational midpoint defined as the PEG200 fraction (% v/v) at which half of the dehydration-responsive folding transition is reached. PEG_T_ values were derived from PEG200 titrations monitored by CD at 265 nm and fitted with normalized sigmoidal transition curves. For example, d(GT)_7.5_ showed a PEG_T_ of 37.6% (Figure 1G). Unlike *T*_m_, which reports thermal unfolding as temperature increases, PEG_T_ reports folding as PEG200 concentration increases.

Two-tetrad LHG4s are often stabilized as stacked two-block assemblies.^54,55,57,58,76^ We therefore tested whether dimeric stacking contributes to the minimal d(GT)_n_ repeat system by adding T_2_ or T_5_ flanking overhangs, which should disfavor end-to-end stacking.^12,77–79^ Addition of terminal thymine flanks reduces the propensity of the sequences to form LHG4. Yet T_5_-d(GT)_7.5_-T_5_ still formed an LHG4-like structure (Figure S7A), indicating that the flanks do not prevent folding. However, compared with unflanked d(GT)_7.5_, the T_5_-flanked construct showed a higher PEG_T_ and a lower *T*_m_ (Figure 1H, I). This is consistent with loss of stabilizing dimeric stacking: the remaining monomeric/intramolecular fold can still form but requires stronger dehydration and is thermally less stable. Native PAGE analysis supported this length-dependent interpretation (Figure S7B). The shortest repeat showed prominent slow-migrating species consistent with dimeric stacking, whereas T_5_ flanking shifted the distribution toward faster-migrating monomeric species. By contrast, T_5_ flanking had smaller effects on the longer repeats, as reflected in their native PAGE profiles and smaller changes in PEG_T_ and *T*_m_ (Figure 1H, I, S7B). Together, these results show that the minimal d(GT)_7.5_ system depends more strongly on dimeric stacking for stabilization than the longer repeats.

The LHG4-like CD signature of d(GT)_7.5_ was retained across the tested micromolar oligonucleotide concentration range (Figure S8). Together with the native PAGE results, this observation is consistent with dimers remaining stable over this range, rather than undergoing appreciable dissociation upon dilution. When T_5_ flanks were added to disfavor dimeric stacking, the PEG_T_ values of d(GT)_7.5_, d(GT)_11.5_, and d(GT)_15.5_ became similar (Figure 1H). This convergence suggests that these repeats have comparable apparent PEG200 requirements when the contribution from dimeric stacking is reduced, whereas the lower PEG_T_ of unflanked d(GT)_7.5_ is consistent with additional stabilization through dimeric stacking.

Together, these results identify d(GT)_n_-derived sequences as dehydration-responsive left-handed G-quadruplexes.

### Mutational analysis reveals sequence constraints on LHG4 folding

We next asked how far this dehydration-responsive folding behavior extends beyond perfect d(GT)_n_ repeats. We constructed and characterized variants from two-tetrad d(GT)_7.5_-derived templates (Table S1). The goal was to define which sequence changes preserve, weaken or abolish drLHG4 folding.

We first tested whether the single-nucleotide T loops in d(GT)_7.5_ could tolerate base substitution. Several loop thymines were substituted with G, C or A. Substitutions are described relative to the parental d(GT)_7.5_ sequence; for example, replacing the first single-nucleotide T loop in d(GT)_7.5_ d-G<u>T</u>GTGTGTGTGTGTG with G gives d-G<u>G</u>GTGTGTGTGTGTG. This substitution retained an LHG4-like CD signature under dehydrating conditions, indicating that the first loop position does not strictly require thymine. More broadly, single T-to-G/A/C substitutions did not systematically abolish left-handed folding or substantially destabilize the structure (Figure S9A, S9B). LHG4 folding tolerated single-base substitutions and multiple T-to-C substitutions, whereas complete replacement of the T loop with A and/or C caused pronounced destabilization, consistent with competing non-LHG4 conformations (Figure S9C, S9D). Thus, LHG4 folding tolerates loop-base substitution at several positions, although this tolerance is sequence- and position-dependent.

We then varied loop length and scaffold continuity through systematic single-thymine insertions (*More*) and thymine deletions (*Less*). For example, *1More4* contains a single inserted thymine at loop position 4 from the d(GT)_7.5_ scaffold, whereas *2Less34* denotes deletion of thymine at the third and fourth loop positions (Figure 2A). Single T insertions were generally destabilizing, consistent with the preference of canonical LHG4s for single-nucleotide loops (Figure S10).^54^ In contrast, single deletions showed strong position dependence. Deletions at loop positions 2, 4 or 6 frequently stabilized the fold, whereas deletions at loop positions 1, 3, 5 or 7 were neutral or destabilizing (Figure 2A, 2B; S11A, S11B). A possible explanation is that even-position deletions preferentially shift the repeat register toward a more stable *Motif1*-like LHG4 arrangement, whereas odd-position deletions may favor a less stable *Motif2*-like arrangement.^55^

**Figure 2.**
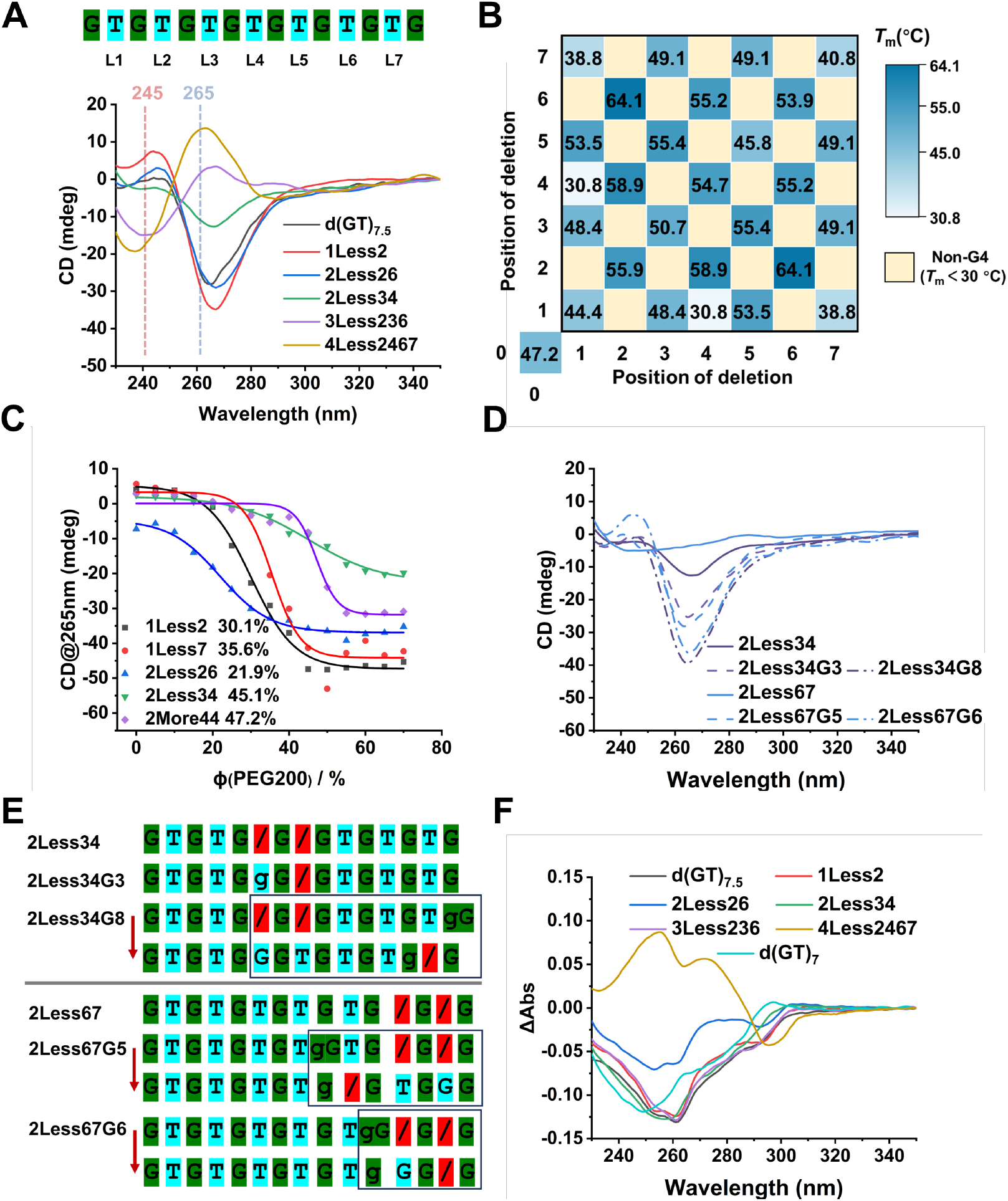
Loop-thymine deletion reshapes the folding landscape of the d(GT)_7.5_ LHG4 scaffold. (**A**) Position code of the d(GT)_7.5_ scaffold and representative CD spectra of selected deletion variants under dehydrating conditions, showing that loop deletions can preserve, weaken, or alter the LHG4-like CD signature. (**B**) Thermal stability matrix for double-position deletion variants. Values indicate *T*_m_ measured at 295 nm in UV-vis absorbance melting, and pale-yellow cells indicate non-G4 variants without an assigned *T*_m_. (**C**) PEG200 titrations monitored by CD at 265 nm, with derived PEG_T_ values. Deletion variants showed distinct dehydration thresholds, indicating position-dependent effects on LHG4 folding. (**D**) CD spectra of selected double-deletion variants and extra-G derivatives, showing that extra G insertion can restore or reshape LHG4-like CD features in otherwise weak deletion scaffolds. (**E**) Register-reassignment model for extra-G rescue variants. Green boxes indicate guanines assigned to the G-quartet core, cyan boxes indicate loop positions, and red boxes containing slashes mark deleted thymine positions. Lowercase ‘g’ identifies the single inserted guanine. Rows connected by red arrows show the same sequence in alternative proposed registers; the arrows indicate reassignment of guanines between quartet and loop positions. Black outlines highlight the sequence regions involved in this reassignment. (F) Thermal difference spectra of representative variants, supporting G4-like folding for variants that retain compatible CD and melting behavior.

We next examined double-deletion variants, where LHG4 folding depended strongly on the relative positions of the two deleted bases. Even-sum deletion mutants often retained LHG4 folding. For example, *2Less26* and *2Less13* remained compatible with left-handed folding, whereas odd-sum variants such as *2Less34* and *2Less25* more frequently disrupted it (Figure 2A, 2B; S11A, S11C, S11D). PEG_T_ measurements showed the same trend (Figure 2C). Thus, deletions influence both local loop architecture and the broader folding behavior of the repeat scaffold.

Introducing a single extra G rescued several deletion variants with weak LHG4-like CD signatures without requiring a specific insertion site (Figure 2D, 2E). Representative examples include insertion adjacent to G3 or G8 in 2Less34, or G5 or G6 in 2Less67. Here, G1-G8 denote the eight guanines of the parental d(GT)_7.5_ scaffold. We interpret this rescue as a register-buffering effect: the additional guanine allows alternative assignments of guanines to the tetrad core. In this model, either the inserted G occupies a loop position, or it enters the core while an original G is reassigned to a loop or bulge position. The extra G therefore allows a more favorable arrangement of quartet-forming guanines and loops, compensating for the disruption caused by thymine deletions.

Triple deletions produced a different outcome. Many triple-deletion variants showed weak CD signals and lacked canonical LHG4 signatures (Figure 2A, S11E, S11F). However, absorbance melting at 295 nm revealed that a subset retained clear thermal transitions and high *T*_m_ values (Table S2, Figure S11A). By contrast, no reliable melting transition at 295 nm could be measured for non-folding controls, such as d(GT)_7_ or odd-pattern double-deletion variants. For *3Less236*, temperature-dependent CD measurements showed that its weak spectral features diminished upon heating (Figure S12). Thermal difference spectra (TDS), an orthogonal UV absorbance readout of nucleic-acid folding, showed G4-like profiles comparable to those of LHG4-forming sequences (Figure 2F).^80^ The combined TDS and melting results therefore support the persistence of thermally stable G4-like states despite the weak CD signals. This weak CD phenotype could reflect conformational heterogeneity or a multimodular structure with compensating contributions from each module to the CD signal.^81–83^ Yet the available data do not distinguish between these possibilities and the structural basis of this behavior warrants further investigation. Further T deletions generated longer contiguous G tracts and disrupted the single-nucleotide loop arrangement characteristic of LHG4s. Consistent with this sequence change, the quadruple-deletion variant *4Less2467* showed an RHG4-like CD spectrum and a G4-like TDS profile (Figure 2A, 2F).

Together, these mutational analyses show that access to LHG4 folding extends beyond the previously defined minimal LHG4 motifs but remains strongly constrained by sequence register.

### Crystallographic characterization of a bulged LHG4 fold

To obtain structural insight into drLHG4 sequences, we attempted to crystallize several of them and successfully solved the structure of *2Less26*, d(GTGGTGTGTGGTG). This sequence folded robustly under dehydrating conditions and showed a relatively low PEG_T_ requirement (21.9%). We obtained two crystal forms in distinct space groups, *P*12_1_1 (0.88 Å resolution, PDB ID 9PLB) and *P*6_5_22 (1.54 Å resolution, PDB ID 9PLC). Data collection and refinement statistics are presented in Table S3. The 0.88 Å resolution of 9PLB places it among the highest-resolution G4 DNA structures reported to date. The overall B-factor for 9PLB structure is significantly lower than that for 9PLC structure (9.5 *vs* 51.8 Å^2^) because of its higher resolution (Figure S13).

Despite different packing arrangements, both structures adopt the same architecture of LHG4 with a thymine bulge (Figure 3A-C). More specifically, all guanines participate in the formation of G-tetrads, T7 forms a bulge within the G6-G8 strand, while T2, T5, T9 and T12 form single-nucleotide loops (note, nucleotide numbering in this case is sequential, with the first guanine labeled as G1 and the last as G13, Figure 3A). The thymine loops fold over each external tetrad of the subunits, forming the thymine caps associated with all reported LHG4 structures.^54-56, 58, 76, 84^ In both crystal forms, the sequence assembles in a dimeric two-subunit LHG4 in which each subunit contributes two stacked G-tetrads, yielding a four-layered structure. In 9PLB, the asymmetric unit (ASU) contains two DNA chains that dimerize directly (Figure 3B), whereas in 9PLC the dimer is formed between one molecule in ASU and a symmetry-generated partner (Figure 3C). Both structures as well as each individual subunit between and within structures are highly similar, with RMSD values between subunits not exceeding 1 Å (Figure S14). These low RMSD values indicate nearly identical folds with greatest differences found in the position of thymine nucleotides in the bulge or loops, which is expected due to their flexibility.

**Figure 3.**
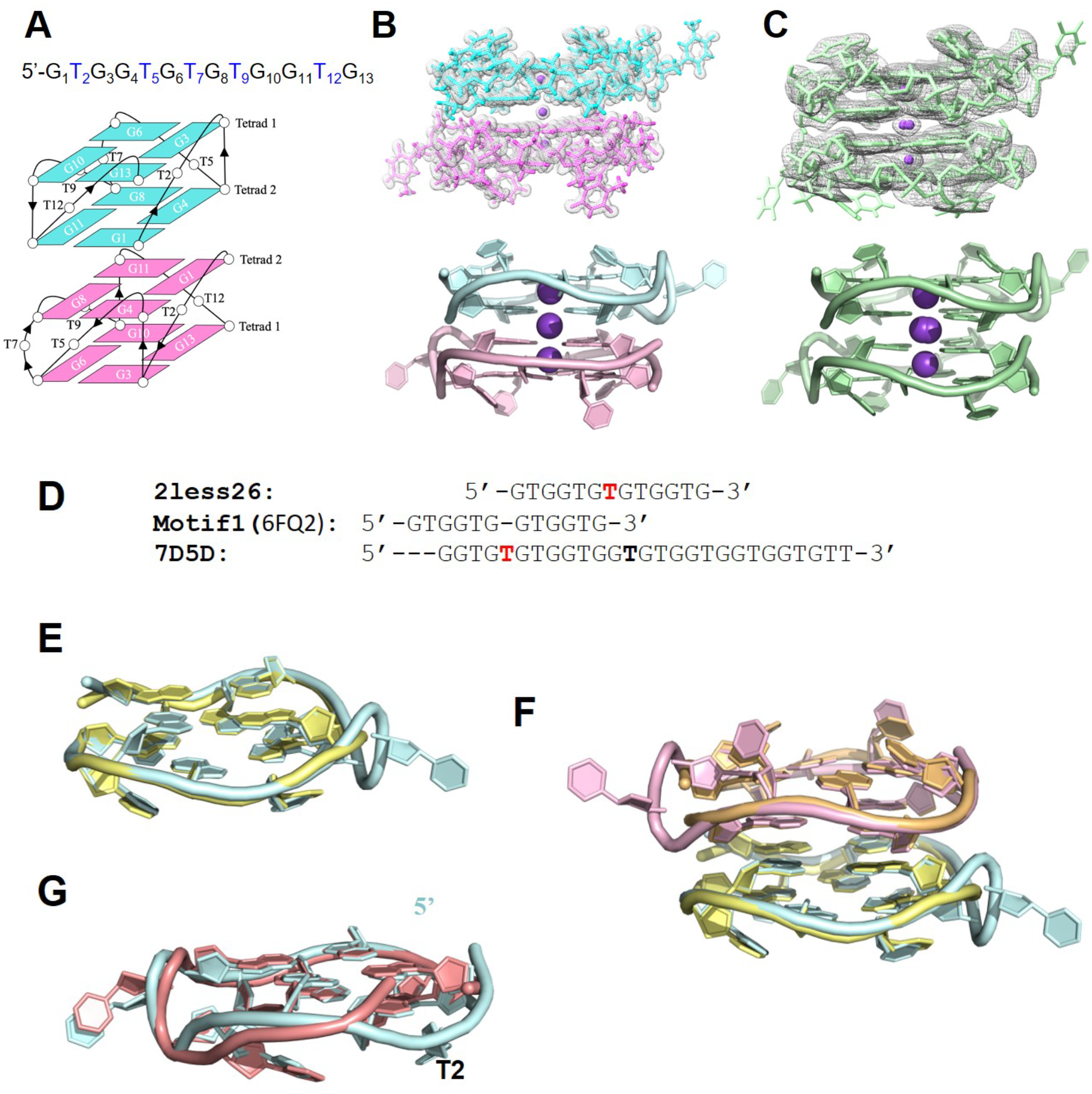
Architecture of *2Less26* structures (PDB ID 9PLB/9PLC) and their comparison to other LHG4 structures. (**A**) DNA sequence and schematic representation of both *2Less26* structures. All bases are numbered, and the arrowed lines indicate backbone progression. (**B**) 9PLB structure with electron density shown at I/σ = 1.0 (top). Cartoon representation of the 9PLB structure (bottom). Chain A is colored cyan, and Chain B is colored salmon. Potassium ions are represented by purple spheres. (**C**) 9PLC structure with electron density shown at I/σ = 1.0 (top). Cartoon representation of the 9PLC structure (bottom). Each monomer is colored pale green. The second molecule is generated by applying symmetry operations. The middle potassium resides on a special position and has been modeled at half occupancy. (**D**) Sequence alignment of *2Less26, Motif1* (PDB ID 6FQ2), and intramolecular LHG4 with bulge (PDB ID 7D5D). Bulge T is in bold red and linker thymine is in bold. The bulge thymine is at a different position in *2Less26* (T7) vs 7D5D (T5) leading to shifted alignment; G1 of *2Less26* aligns with G12 of 7D5D; T2 in *2Less26* does not have an equivalent nucleotide in 7D5D. (**E**) Overlay of Chain A of 9PLB (light blue) with chain A of *Motif1* (PDB ID 6FQ2, yellow). (**F**) Overlay of chain A/B (blue/pink) of 9PLB with chain A/B (yellow and orange) of *Motif1* (PDB ID 6FQ2). (**G**) Overlay of 9PLB (light blue) and the bulge-containing subunit of LHG4 with PDB ID 7D5D (salmon).

G-tetrads in both structures are relatively planar, with out-of-plane deviations (*D*_OOP_) of 1.0-1.3 Å, comparable to that observed in other LHG4s (Table S4). The overall architecture of LHG4 is further stabilized by three potassium ions per dimer, one of which resides at the dimer interface (Figure 3B-C). Potassium coordination distances are within the expected range for G4 central channels (Figure S15). ^54-56, 58, 76, 84, 85^ Groove width measurements are consistent with a parallel topology with all medium grooves measuring 14.5-15.4 Å (Table S5) in agreement with the 15.1 ± 0.7 Å reported for 333 intramolecular G4 structures.^85^ The backbone follows the characteristic left-handed strand progression with the negative helical twist between two tetrads of about -26° and between subunits (in 9PLB only) of -11.0° (Table S6). The observed values of helical twists are consistent with those reported by our laboratory for the left-handed subunit of the G4 structure from SLC2A1 promoter.^58^ Torsion angle analysis of both structures indicates no outliers and further supports the high quality of both models (Figure S16).

The repetitive nature of the LHG4-forming sequences (*Motif1* and *Motif2*) and their nearly invariant fold allow for an effective alignment of LHG4 structures (Figure 3D-F). Excluding the T7 bulge, *2Less26* is nearly identical to the *Motif1* monomer (PDB ID 6FQ2), both in sequence and in structure (RMSD = 0.68 Å) (Figure 3E). The overall organization of the *2Less26* and *Motif1* dimers is also very similar (Figure 3F), maintaining the same positions of G-tetrads and thymine caps (RMSD = 0.70 Å, excluding phosphate backbone) but differing in the rotational relationship between the two monomers, and, thus, the organization of their backbones (i.e., locations of the interdomain “break”). Preservation of the *Motif1*-like core while accommodating an additional thymine as a bulge provides a structural example of the sequence tolerance observed in the mutational analysis. The stacked two-block architecture is consistent with the stabilizing role proposed for dimerization in the minimal-repeat system. Previous studies have shown that an LHG4 motif can also promote left-handed folding in a compatible adjacent sequence, highlighting the ability of interblock interactions to influence folding preferences.^54-56, 84^

In earlier work from the Phan lab, sequences designed to accommodate one to four bulges were investigated by linking them to an intact *Motif1* block, which promoted and stabilized left-handed folding in the adjacent G-rich block. Structures containing one to three bulges were determined by NMR or X-ray crystallography. Our *2Less26* structure is similar to the structure with a single bulge (PDB ID 7D5D).^84^ After excluding T2 of *2Less26*, which has no equivalent in 7D5D, the two structures aligned with an RMSD of 0.95 Å (Figure 3G). The four-bulge construct remained unstructured under the conditions examined.^84^ Without the appended *Motif1* block, the corresponding sequences did not independently form well-defined LHG4s. Interestingly their 4-bulge block sequence is identical to our d(GT)_7.5_. However, they only explored dilute buffer conditions. We can generalize our structural finding to all studied d(GT)_n_ sequences (n > 7) and their variants: under specific environmental conditions, the sequences can adopt the LHG4 fold by utilizing their guanines for G-tetrad formation, certain thymines as 1-nt loops that stabilize parallel fold and pushing unused thymines outward, leading to bulges. Importantly, the Phan lab showed that bulges in LHG4 are not as destabilizing as they are in RHG4.^84^

### Hydration sphere and PEG200 in *2Less26* crystals

Because of their importance to this work, we focus here on hydration sphere and PEG200 in *2Less26* structures. The intermolecular contacts as well as the presence of other molecules and their roles are described in detail in the SI section (Figure S17). The 0.88 Å structure of 9PLB enabled unusually detailed analysis of hydration around the left-handed fold. The ASU of 9PLB contains 132 water molecules, which form an extensive network around the exterior of the LHG4 dimer. This water network displays similar features to that of the first discovered LHG4 (PDB ID: 4U5M).^56,68^ Most water molecules are concentrated near the circular groove between two subunits (Figure 4A). Only a few interior waters in this region interact directly with guanines via their N2 and N3 (shown in green in Figure 4B, 4C), while the rest (shown in magenta) interact with the sugar-phosphate backbone. These exterior waters form a near-continuous “wheel” around this circular groove (Figure 4C). The water wheel reveals an organized hydration network around the folded dimer. Its spatial arrangement suggests that solvent-dependent stability may involve local reorganization of hydration, rather than being determined by bulk water activity alone.

**Figure 4.**
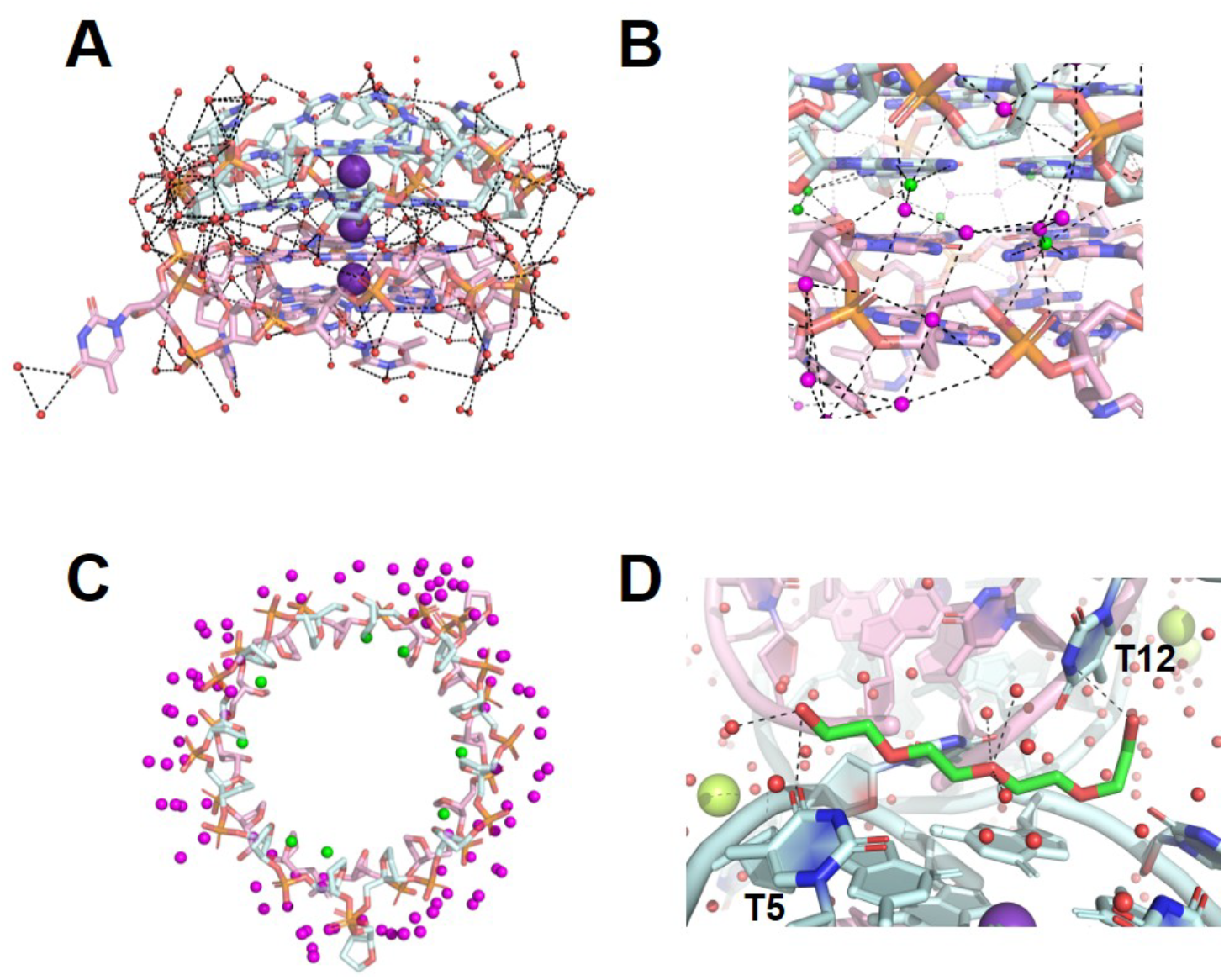
Hydration sphere and PEG200 in *2Less26* crystals. (**A**) An entire water network. DNA chains are colored as in Figure 3. Waters are shown as red spheres. Na^+^ and PEG are omitted for clarity. Hydrogen bonds are shown as black dashed lines and are only drawn for distances < 3.2 Å. (**B**) Groove water network. Interior waters (green) hydrogen bond to N2 and N3 of guanines, while exterior waters (magenta) hydrogen bond to other waters or the phosphate-sugar backbone. **(C**) View of the “water wheel” looking down the ion channel. Only the sugar-phosphate backbone of DNA is shown. (**D**) Interactions involving PEG200 molecule in 9PLB structure. PEG is colored by atom: carbons are forest green while oxygens are red.

Our biophysical work demonstrates that PEG200 effectively promoted LHG4 folding in our solution experiments. Excitingly, both structures crystallized with PEG200 which occupies a pocket between thymine-cap regions. In the 9PLB structure we can see the entirety of the PEG200 molecule. Its terminal hydroxyl groups contact T2, T5 and T12 loop thymines in 9PLC, and T5 and T12 in 9PLB (Figure 4D, S17E). The exact hydrogen-bonding geometry should be interpreted cautiously, because the terminal hydroxyl groups of PEG200 were modeled with some ambiguity. Nevertheless, the PEG200 density and position are clear in both structures. Thus, it is possible that the role of PEG200 is to locally stabilize the thymine-cap region, as analogous sites in related LHG4 structures are occupied by PEG oligomers or the polyamine spermine (in molecules with PDB ID 7D5D, 7D5E, 8TAA, 9CIY, 9O70, 9MXO, 9YYP, 11XE). However, these crystallographic contacts do not by themselves establish whether PEG200 promotes LHG4 formation through direct binding, dehydration, or both, because chemically distinct dehydrating agents, including ethylene glycol, DMSO and acetonitrile, also induce LHG4 folding. These observations suggest that PEG200 may combine changes in the bulk solvent environment with additional local stabilization of the thymine-cap region. Together, our structures reveal a distinct solvent organization that helps explain the strong coupling between LHG4 folding and dehydration.

### Molecular binders induce and stabilize drLHG4 states

Previous studies have shown that small-molecule ligands can promote G4 formation without requiring an initially stable, folded G4 population.^86^ Building on this precedent and our observation that solvent conditions promote LHG4 folding in repeat-derived sequences, we next asked whether selected G4-binding peptides or ligands could promote drLHG4 formation in the absence of bulk dehydrating cosolvents. We hypothesized that binding could favor folded conformations, potentially aided by interfacial hydration changes, as molecular recognition often involves reorganization or displacement of interfacial water.^87–88^

We screened a panel of published G4-binding peptides and protein-derived fragments (Table S7, Figure S18), including the RHAU-derived peptides RHAU23 and G4P,^29,89^ CIRBP-peptide12,^90^ WRN-RQC,^91^ NPM-C70,^92^ and KR12C^93^ against drLHG4-forming sequences. Only G4P, and to a lesser extent RHAU23, reproducibly induced drLHG4 in the absence of cosolvents (Figure 5A). Fluorescence anisotropy was used to estimate apparent binding affinities. The data showed strong sequence and peptide dependence. Specifically, d(GT)_7.5_ bound tightly to G4P (with an apparent *K*_d_ of 14.4 ± 4.3 nM), much more weakly to RHAU23 (*K*_d_ = 1.05 ± 0.50 μM), and did not bind other peptides (Figure 5B). We also examined *3Less236* to test whether peptide binding extended to a variant lacking a canonical LHG4-like CD signature but retaining a clear thermal transition and a G4-like TDS profile. A similar binding trend was observed for this variant, with apparent *K*_d_ values of 12.8 ± 4.1 nM for G4P and 0.68 ± 0.27 μM for RHAU23 (Figure 5C). In contrast, no peptides yielded measurable binding to d(GT)_7_ (Figure S19). These results indicate that drLHG4 induction is not a general property of G4-binding peptides but is preferentially associated with RHAU-derived tetrad-recognition modules. G4P contains tandem RHAU-derived modules, which may provide a multivalent binding advantage allowing more efficient capture and stabilization of inducible drLHG4 states than the single RHAU23 peptide.^89^

**Figure 5.**
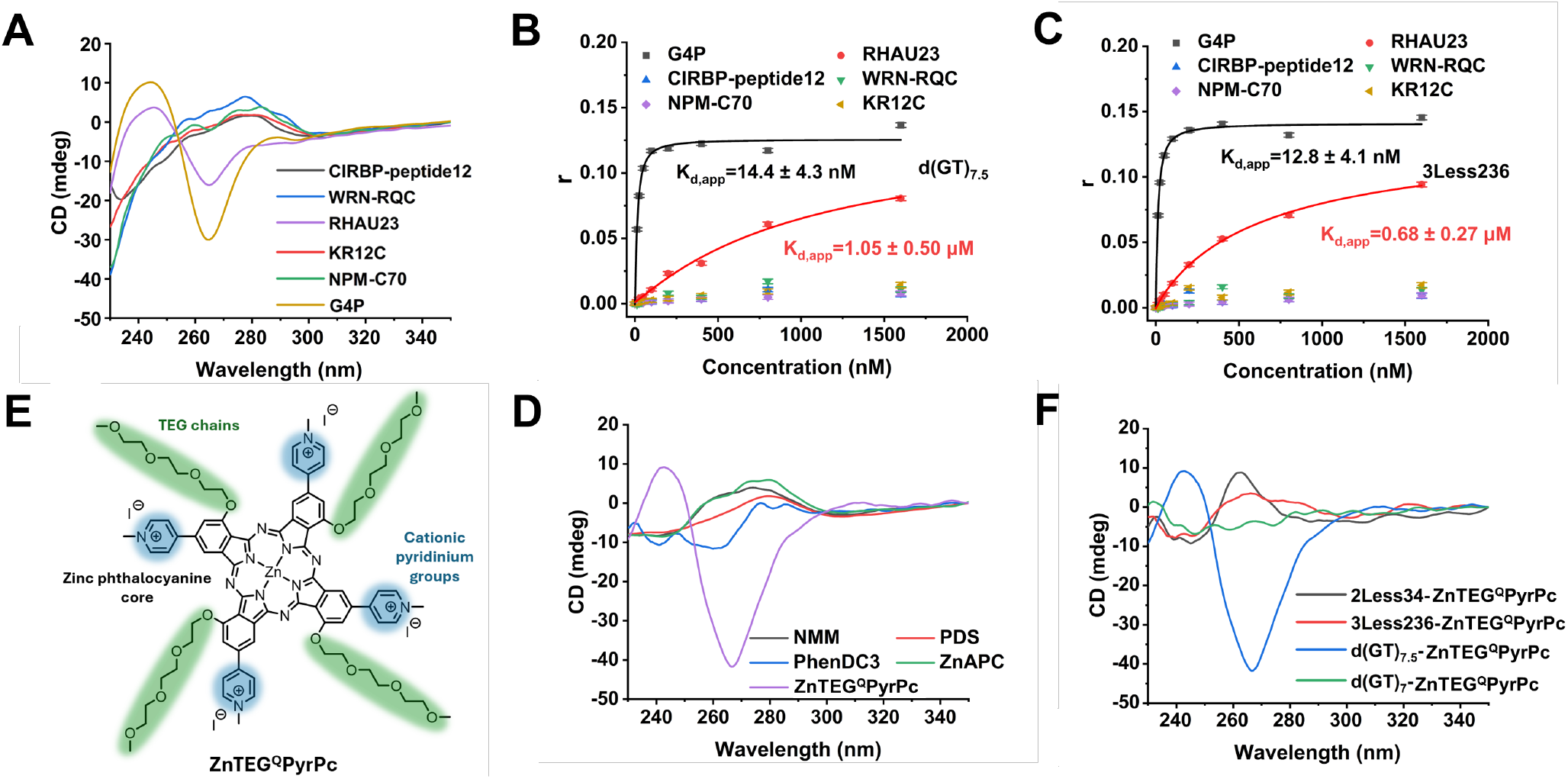
Peptides and small-molecule ligands induce drLHG4 folding. (**A**) CD spectra of 5 μM d(GT)_7.5_ in the presence of 10 μM G4-binding peptides and protein-derived fragments. (B, C) Fluorescence anisotropy titrations of peptide binding to d(GT)_7.5_ (**B**) and *3Less236* (**C**). Curves represent one-site apparent binding isotherm fits, and apparent *K*_d_ values are indicated. Data are presented as mean ± SD from n = 3 independent measurements. (**D**) CD spectra of 5 μM d(GT)_7.5_ in the presence of 12.5 μM of conventional G4 ligands or ZnTEG^Q^PyrPc. (**E**) Chemical structure of ZnTEG^Q^PyrPc, highlighting the zinc phthalocyanine core, cationic pyridinium groups, and TEG (triethylene glycol methyl ether) tails. (**F**) CD spectra of 5 μM selected sequences in the presence of 12.5 μM ZnTEG^Q^PyrPc.

We then tested classical small-molecule G4 binders which included NMM,^94^ pyridostatin (PDS),^95^ zinc phthalocyanine tetrasulfonic acid (ZnAPC),^96^ and PhenDC3.^97^ None induced drLHG4 formation (Figure 5D). Fortuitously, we identified the phthalocyanine ZnTEG^Q^PyrPc; this compound exhibits high binding affinity for the d(GT)_10_ sequence but relatively weak binding to RHG4 structures, including d(G_4_T_4_G_4_), d(GG)_10_, and the human telomere sequence.^98^ ZnTEG^Q^PyrPc is a zinc phthalocyanine complex with four triethylene glycol methyl ether (TEG) substituents (Figure 5E), which exists in an aggregated state in solution. Binding to d(GT)_10_ DNA disrupted the aggregates as was observed via spectral changes in the absorption spectra and emission enhancement of the monomeric phthalocyanine absorption Q-band.^98-100^ In addition, it led to significant changes in the CD spectra of d(GT)_10_ with a negative band at 266 nm and positive band at 244 nm, reminiscent of LHG4 topology.^98^ This selective binding to d(GT)_10_ suggested that ZnTEG^Q^PyrPc might engage other GT-repeat sequences capable of adopting drLHG4-like states. Thus, we carried out detailed investigations to examine how generalizable this interaction is. We discovered that 2.5 equivalents of ZnTEG^Q^PyrPc can convert d(GT)_7.5_ into drLHG4 in the absence of PEG200, while d(GT)_7_ was left unchanged (Figure 5F). We also monitored aggregation state of ZnTEG^Q^PyrPc upon addition of d(GT)_7.5_ and d(GT)_7_ and only observed significant changes in the UV-visible absorption and fluorescence emission spectra upon addition of the former sequence, while d(GT)_7_ induced only minor spectral changes, likely reflecting electrostatic interactions (Figure S20). Overall, we conclude that ZnTEG^Q^PyrPc can induce drLHG4s via formation of DNA-bound phthalocyanine monomers in solution. Of note is the similarity in length of TEG (three ethylene glycol units) and PEG200 (four units) that also induces drLHG4 formation. Therefore, the unique and contrasting behavior of ZnTEG^Q^PyrPc to other small molecules, including ZnAPC, may be attributed to the presence of the TEG chains and the ability of the bound ZnTEG^Q^PyrPc to position these chains on the G4 to generate positive interactions and a local dehydrating environment. This is likely facilitated by terminal binding of the ZnTEG^Q^PyrPc to LHG4. A more recent study has investigated the selective binding of this same ligand to i-motif structures where the TEG substituents may also be highly relevant for the binding.^99^

Additional sequence variants also promoted disaggregation of ZnTEG^Q^PyrPc aggregates (Figure S21), although their CD spectra did not develop canonical LHG4-like profiles (Figure 5F). We next compared the sequence-dependent fluorescence response of ZnTEG^Q^PyrPc upon DNA titration (Figure S22). Because ZnTEG^Q^PyrPc is aggregated under these conditions and DNA binding is coupled to aggregate disruption and monomer stabilization, the fitted values are treated as apparent half-maximal response concentrations rather than microscopic 1:1 dissociation constants. Titration with d(GT)_7.5_ yielded a *K*_1/2,app_ value of 56.0 nM, compared with 69.2 nM for *3Less236* and 487 nM for *2Less34*. In contrast, d(GT)_7_ showed no measurable saturation within the tested concentration range and produced a much weaker fluorescence response.

## Discussion

LHG4s have so far been viewed as a rare structural class confined to closely related sequence motifs. Our results challenge this notion. Simple d(GT)_n_-rich repeats, together with their variants, can access LHG4 states when the hydration environment or local molecular interface favors their formation. These findings expand the LHG4 sequence repertoire beyond established motifs and identify dehydration as a condition that opens access to this broader folding landscape.

The sequences described here are not simply “G4-forming” or “non-G4-forming” in an absolute sense. Instead, their folding depends on repeat length, K^+^ concentration, conditions that promote productive DNA dehydration and, in minimal systems, dimeric stacking. Bulk water activity alone does not predict productive LHG4 folding: chemically distinct cosolvents with similar water-activity ranges produced different folding outcomes. The Ficoll 70 and high-molecular-weight PEG results further separate productive dehydration-responsive folding from generic excluded-volume crowding.^39–41,101^ We therefore interpret d(GT)_n_-derived LHG4 formation as a locally gated dehydration process that requires remodeling of the hydration shell around nascent tetrad-, loop- and bulge-rich interfaces. The environment thus determines not only the stability of a folded structure, but also which sequences can access it.

Under standard aqueous conditions, substitutions and insertions in *Motif1*-derived sequences weakened or disrupted the characteristic LHG4-like CD signature (Figure S23). Previous work on two-block LHG4s showed that an intact *Motif1* block stabilizes a mutated neighboring block, whereas mutations in both blocks compromise LHG4 formation, highlighting the stabilizing role of a preserved motif under these conditions.^54^ In contrast, under dehydrating conditions, our mutational analysis reveals a broader range of LHG4-compatible sequences. Loop substitutions can be tolerated, deletion effects depend on the positions and combinations of deleted thymines, and an extra guanine can rescue otherwise inactive variants. This expanded tolerance is nevertheless not a consequence of G-richness alone: single-nucleotide loops remain strongly favored in the two-tetrad system, and the arrangement of guanines and loops determines the folding outcome. Some variants retain canonical LHG4 signatures, whereas others form thermally stable but CD-weak G4-like states that will require further investigation.

The crystal structures of *2Less26* provide a structural explanation for this sequence tolerance: a repeat-derived sequence preserves the core LHG4 fold by moving a thymine nucleotide into a bulge. Previous work established bulged LHG4 structures by linking the bulge-forming sequence to an intact *Motif1* block that promoted left-handed folding in the adjacent block.^84^ Here, *2Less26* adopts a bulged LHG4 architecture without an appended intact *Motif1* block, and two crystal forms converge on the same overall architecture. Together with the solvent-dependent folding observed in solution, these structures suggest that dehydrating conditions can make bulged LHG4 formation more favorable relative to unfolded or competing conformations. The 0.88 Å 9PLB structure further reveals an extended water network around the circular groove. We hypothesize that reorganization of water at this interface contributes to the dehydration dependence observed in solution. The observed PEG200 molecule may additionally stabilize the thymine-cap arrangement. However, induction by chemically distinct dehydrating agents argues against a purely PEG-specific ligand effect: local stabilization by PEG200 complements a broader dehydration-responsive folding mechanism rather than being the sole origin of LHG4 induction.

Molecular recognition provides another route into this folding landscape without bulk dehydrating cosolvents. Only a subset of the tested G4 binders promoted drLHG4 folding, distinguishing folding induction from binding alone. Here, induction refers to the appearance of an LHG4-like CD signature upon binder addition. We hypothesize that certain binders preferentially stabilize drLHG4-compatible intermediates or folded conformations, thereby shifting the folding equilibrium toward drLHG4. RHAU23 and G4P may act through geometrically matched contacts with exposed tetrad surfaces, with the stronger inducing activity of G4P reflecting more effective stabilization by its tandem RHAU-derived modules. The small-molecule binding results reinforce this distinction: conventional G4 ligands did not induce comparable LHG4-like CD signatures, whereas ZnTEG^Q^PyrPc was effective. An approximately 1:1 ligand ratio was sufficient to promote LHG4 formation, corresponding to four ligand-borne TEG chains added per d(GT)_7.5_ molecule, in contrast to the large excess of bulk PEG200 required. We hypothesize that the phthalocyanine core positions these chains close to the DNA surface, combining G4 stabilization with local hydration remodeling. This proposed mechanism remains to be tested; the present results demonstrate binder-dependent induction but do not establish the underlying changes in interfacial hydration.

Our findings have implications beyond the sequences examined here. d(GT)_n_ and related tracts may not be suitable single-stranded repeat controls, as they can adopt G4-like structures under activating conditions that standard aqueous assays fail to capture. Protein binding and other cellular processes that alter local nucleic-acid environments could provide routes into this dehydration-responsive state space.^102–104^ Budding yeast telomeres offer a concrete example: their heterogeneous TG_1–3_ tracts differ from regular mammalian telomeric repeats,^105,106^ yet a representative truncated yeast telomeric sequence formed a stable LHG4 structure under dehydrating conditions (Figure S24). Whether such states are accessed in cells remains to be established. More broadly, the question shifts from which G-rich sequences form G4s in standard buffers to which remain unable to do so across the diverse local environments that biological systems may provide. Failure to fold under conventional in vitro conditions may reflect the conditions tested rather than an intrinsic limitation of the sequence.

## Conclusions

In summary, this work expands LHG4s from a narrow motif-defined class to a dehydration-responsive family of left-handed G4 states. d(GT)_n_ repeats and their variants reveal position-dependent sequence requirements, while the *2Less26* crystal structures provide direct evidence for a repeat-derived bulged LHG4 architecture and a distinctive hydration network. The unique ability of selected peptides and ZnTEG^Q^PyrPc to fold LHG4s shows that these states can be induced by local binding interfaces and can direct the future development of LHG4-specific binders. These findings define a framework for studying condition-accessible G4 structures beyond those populated in standard aqueous buffers.

## Acronyms

G4: G-quadruplex
LHG4 and RHG4: left-handed and right-handed G-quadruplex, respectively
drLHG4: dehydration-responsive left-handed G-quadruplex
CD: circular dichroism
TDS: thermal difference spectrum
*T*_m_: melting temperature
PEG: polyethylene glycol
PEG_T_: PEG200 fraction (% v/v) at the midpoint of the dehydration-responsive folding transition
ZnTEG^Q^PyrPc: cationic zinc(II) phthalocyanine derivative with four triethylene glycol methyl ether tails in the α positions and four N-methylpyridinium groups in β positions.

## Supplementary Data

Supplementary Data are available online.

## Acknowledgments

We thank Anne Cucchiarini and Lionel Guittat (LOB, Palaiseau) for technical assistance and helpful advice.

## Author Contributions

Y.C. conceived the project and carried out most of the experiments; E.R.W. performed ZnTEG^Q^PyrPc spectroscopic assays; J.W. assisted with peptide synthesis and screening; A.H. performed crystal screening and solved the structure; E.X. analyzed the structure. Y.C., J.W. and J.-L.M. analyzed the data and interpreted the results. Y.C., L.A.Y. and J.-L.M. wrote the manuscript, with contributions from J.G., E.R.W., R.M.E. and S.J.Q., who also critically reviewed and revised it. R.M.E. developed ZnTEG^Q^PyrPc along with E.R.W. and S.J.Q. J.-L.M. provided academic mentorship to Y.C. All authors critically read the final manuscript.

## Funding

This work was supported by the *Fondation de l’École polytechnique* (project Pathogens), Centre National de la Recherche Scientifique (CNRS) and Institut National de la Santé et de la Recherche Médicale (Inserm). Y.C. is the recipient of a China Scholarship Council PhD grant (202308310081). L.A.Y. acknowledges support from Swarthmore College research and start up funds. The Center for Bio-Molecular Structure (CBMS) is primarily supported by the NIH-NIGMS through a Center Core P30 Grant (P30GM133893), and by the DOE Office of Biological and Environmental Research (KP1607011). NSLS-II is a U.S. DOE Office of Science User Facility operated under Contract No. DE-SC0012704. This publication resulted from the data collected using the beamtime obtained through NECAT BAG proposal #317877. E.R.W. acknowledges financial support from the Research Ireland (GOIPG/2022/1362). S.J.Q. acknowledges financial support from the Research Ireland Photogene project (21/FFP-P/10126). R.M.E. acknowledges financial support from the University of Strathclyde for a Chancellor’s Fellowship and the South African Medical Research Council with funds received from the South African National Department of Health and the UKRI Medical Research Council via the UK Government’s International Science Partnerships Fund.

## Conflict of interest

None declared.

## Entry for the Table of Contents

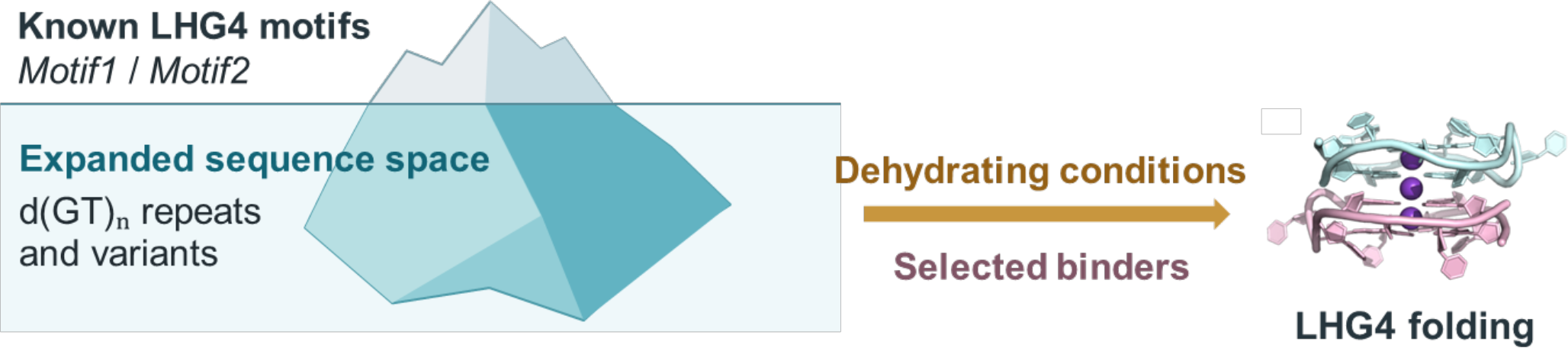

Known G4 motifs are only the tip of the iceberg. Dehydrating conditions and selected molecular binders unlock left-handed G-quadruplex folding in simple d(GT)_n_ repeats and their variants. Crystal structures reveal how bulges preserve the left-handed core, extending this fold beyond its established sequence boundaries.

## Notes

### Competing Interest Statement

The authors have declared no competing interest.

### Summary of Updates

The title and abstract copied / pasted in the BioRXiv server were NOT the right ones - Very sorry for this mistake on our side. The new tittle and abstract are identical to the ones in the pdf. If possible do not publish the version with the incorrect title and abstract

