## Supplementary information for "Dehydration-Responsive Left-Handed G-Quadruplexes Expand the Sequence Repertoire"

### Supporting Information

#### Materials and Methods

The synthesis of the zinc phthalocyanine derivative $ZnTEG^{Q}PyrPc$ was reported previously.[1]

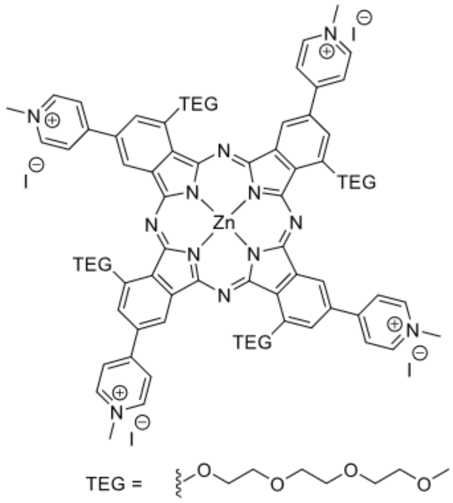


#### Oligonucleotides, ligands, and peptides

All sequences were purchased lyophilized from Eurogentec (Belgium) with RP-cartridge purification and resuspended in a proper volume of ultrapure water to a final concentration of 100 μM. Sequences are provided in Table S1. Concentrations were calculated using the extinction coefficients provided by the manufacturer. All sequences were annealed at 95 °C in an appropriate buffer containing 30 mM Tris-HCl (pH 7.4) for 5 min and cooled down slowly until reaching room temperature. Unless otherwise specified, all spectral assays were performed with 5 μM DNA in 30 mM Tris-HCl, pH 7.4, 50 mM $K^+$, with or without 75% PEG200.

All peptides used in this study are listed in Table S7. RHAU23, WRN-RQC, CIRBP-peptide12 and KR12C were chemically synthesized by Sangon Co., Ltd. (China), whereas G4P and NPM-C70 were recombinantly expressed in *Escherichia coli* in our laboratory. The G4P expression plasmid was kindly provided by Prof. Zheng Tan's laboratory. For NPM-C70, mRNA was extracted from A549 cells using the Dynabeads™ mRNA Direct Kit (Thermo Fisher Scientific) and reverse-transcribed using the RevertAid First Strand cDNA Synthesis Kit (Thermo Fisher Scientific). The sequence encoding NPM-C70 was PCR-amplified using high-fidelity DNA polymerase and primers containing EcoRI and HindIII restriction sites. The amplified product was gel-purified, and both the insert and pET-28a vector were digested with EcoRI and HindIII at 37 °C for 1 h. Following purification, the insert and vector were ligated using T4 DNA ligase at 22 °C for 10 min. The ligation mixture was introduced into competent *E. coli* DH5α cells by heat-shock transformation. Transformants were selected on kanamycin-containing LB agar, and positive clones were verified by sequencing.

G4P and NPM-C70 were expressed using their respective plasmids in *E. coli* Rosetta (DE3) cells. Cultures were grown in LB medium containing 30 μg $mL^{-1}$ kanamycin and 20 μg $mL^{-1}$ chloramphenicol at 37 °C until $OD_{600}$ reached 0.7. Expression was induced with 0.5 mM IPTG at 16 °C overnight. Harvested cells were resuspended in 30 mM HEPES, pH 7.4, containing 150 mM NaCl, 1 mg $mL^{-1}$ lysozyme and protease inhibitors. Following incubation on ice for 20 min, cells were disrupted by eight sonication cycles, each comprising a 20-s pulse followed by a 60-s interval. Lysates were clarified by centrifugation at 4 °C.

Each His-tagged protein was purified using a 5-mL HisTrap Ni-affinity column on an ÄKTA pure system, with an imidazole gradient from 20 to 500 mM. Purified proteins were desalted using an Econo-Pac 10DG column (Bio-Rad) and concentrated using Amicon Ultra centrifugal filters. Glycerol was added to a final concentration of 10%, and samples were stored at −20 °C. The His tags were retained in the proteins used for subsequent experiments. Purification fractions were analyzed by Coomassie Brilliant Blue-stained SDS-PAGE and Western blotting using an anti-His antibody (Figure S25).

**UV-visible absorption spectroscopy**

UV-visible absorption spectra were recorded on an Agilent Cary 3500 spectrometer equipped with a Peltier temperature controller (temperature accuracy ±0.5 °C). Spectra were acquired from 220 to 330 nm. UV melting tests were monitored at 295 nm. Samples were heated from 5 to 95 °C at 0.2 °C/min, with 3 min at each temperature point.

**Thermal difference spectra**

Thermal difference spectra (TDS) were calculated by subtracting UV-vis spectra collected at 20 °C from spectra collected at 90 °C. Samples were equilibrated for 5 min at 20 °C and for 10 min at 90 °C before acquisition. The lower and upper temperatures were chosen to approximate predominantly folded and predominantly unfolded states, respectively.

TDS was calculated as: $TDS(\lambda) = A(90\ °C)(\lambda) - A(20\ °C)(\lambda)$. TDS was used as an orthogonal indicator of G4 formation, particularly for sequences that did not display canonical left-handed CD signatures.

**Circular dichroism spectroscopy**

CD spectra were recorded on a Jasco J-1500 circular dichroism spectropolarimeter equipped with a Peltier temperature controller (temperature accuracy ±0.5 °C). Unless otherwise stated, spectra were collected at 5 °C in 1 cm quartz cuvettes from 220 to 350 nm. Three scans were collected and averaged for each sample. For ion-dependence, PEG-dependence and binder-induction experiments, spectra were collected under the indicated conditions using otherwise identical acquisition parameters. Molar circular dichroism was expressed as Δε and calculated from the measured ellipticity according to $\Delta\varepsilon = \theta / (32980 \times c \times l)$, where θ is the ellipticity in millidegrees (mdeg), c is the DNA concentration in mol/L, and l is the optical path length in cm.

**CD thermal melting**

CD melting tests were monitored at the wavelength corresponding to the dominant positive band of the structure. Samples were heated from 5 to 95 °C at 1.0 °C/min, with 3 min at each temperature point.

**Water activity measurements**

Water activity ($a_w$) was measured at 25 °C using an AQUALAB TDL 2 water activity meter (Addium Inc.) equipped with a tunable diode laser sensor. PEG200, PEG600, DMSO, acetonitrile and glycerol concentrations were expressed as % (v/v), whereas PEG8000 concentrations were expressed in mg/mL. Solutions were mixed until homogeneous and maintained in sealed containers before measurement. Instrument performance was verified using certified water-activity standards covering the measurement range, with calibration adjusted as necessary. Each sample was transferred to a clean sample cup in sufficient volume to cover the base, and the chamber was closed promptly to minimize evaporation. Final readings were recorded after the instrument indicated measurement completion. Three separately loaded aliquots of each solution were analyzed, and water activity was reported as the mean ± SD and plotted against solute concentration.

**Determination of $PEG_T$**

$PEG_T$ was defined as the PEG200 fraction at which half of the dehydration-induced folding transition is reached. To determine $PEG_T$, CD spectra were collected across a PEG200 titration series under otherwise identical buffer conditions. $PEG_T$ was extracted as the PEG200 fraction corresponding to half-maximal transition.

**Native PAGE analysis**

DNA samples (5 μM) were annealed in folding buffer containing 50% PEG200 and 100 mM KCl before loading. Samples were mixed with native loading buffer containing the same PEG200 concentration to minimize perturbation of the folded state during loading. The samples were resolved on a 15% non-denaturing polyacrylamide gel containing 100 mM KCl, 40% (w/v) PEG200. Electrophoresis was performed at 4 °C at 80 V. The gel was stained with SYBR Gold and imaged using a gel documentation system. Band assignments were made from relative migration: slower-migrating bands were assigned as dimeric or stacked species, whereas faster-migrating bands were assigned as monomeric species.

**NMR spectroscopy**

NMR experiments were carried out on a 600 MHz Bruker Avance III spectrometer (Bruker, Germany). For NMR titrations, 200 µM DNA samples were prepared in the buffer containing 50% $H_2O$ and 50% $d_6$-DMSO, ns = 512 cycles (30 min).

**Crystallization**

Crystallization was achieved at 12 °C using the hanging-drop vapor diffusion method. Samples were prepared by annealing 1.5 mM DNA in 50 mM KCl, 10 mM Bis-Tris, pH 7.0, and 30% PEG200. Screening trays were set with the SPT Labtech Mosquito LV Crystal robot equipped with a humidity chamber using 100 µL per well Helix (Molecular Dimensions) and Natrix (Hampton Research) conditions. The drops contained 0.2 µL of DNA sample and 0.2 µL of crystallization condition. An initial hit was found in Natrix 2-16, consisting of 80 mM $SrCl_2$, 40 mM sodium cacodylate, pH 6.0, 35% (w/v) 2-methyl-2,4-pentanediol (MPD), and 12 mM spermine tetrahydrochloride. Optimization trays were set manually using 400 µL crystallization condition per well and optimizing MPD (31-41% *w*/*v*) and $SrCl_2$ (40 mM to 140 mM). The drops contained 1 µL of DNA sample and 1 µL of crystallization condition. Large, flat hexagonal diffraction-quality crystals grew within 2 weeks and yielded the structure with the space group of $P6_522$ (PDB ID 9PLC). We have also swapped $SrCl_2$ for $CaCl_2$ and increased the buffer pH from 6.0 to 6.5. The most successful condition contained 80 mM $CaCl_2$, 40 mM sodium cacodylate, pH 6.5, 37% (w/v) MPD, and 12 mM spermine tetrahydrochloride. Flat, rhomboidal diffraction-quality crystals grew within 24 hours of setting and yielded the structure with the space group of $P12_11$ (PDB ID 9PLB). Crystals were flash frozen in liquid nitrogen without additional cryoprotection due to the high MPD concentration.

Data were collected at the Advanced Photon Source (APS) 24 ID-E synchrotron facility to a maximum resolution of 1.54 Å for 9PLC and the National Synchrotron Light Source II (NSLS II) 17-ID-2 (FMX) to a maximum resolution of 0.88 Å for 9PLB. Raw diffraction data were processed using RAPD at the APS and AutoPROC at the BNL.[2] The structure was solved via molecular replacement (MR) using Phenix.Phaser (version 1.21.2_5419).[3] For MR we used one molecule from the left-handed DNA G-quadruplex structure with PDB ID 6FQ2. Extensive manual model building cycles were performed using Coot 0.8.8 followed by Phenix.Refine.[4-5]

The asymmetric unit of 9PLB contains two DNA chains (with hydrogen atoms added at idealized positions), 132 water molecules, three potassium ions, two sodium ions, and one PEG200. The high resolution of this structure allowed us to visualize PEG200 as $H(\text{-O-}CH_2\text{-}CH_2\text{-})_4OH$, containing four repeats. This PEG indeed should be the most prevalent species in the mixture labeled as PEG200. Our choice of $Na^+$ but not $Ca^{2+}$ relies on marginally better $R_{work}/R_{free}$ values (~0.1% difference) but the structural parameters for both metals are rather similar and CheckMyMetal site[6] equally accepts both. Notably, refining occupancies for $Ca^{2+}$ yielded values between 0.6 and 0.7, suggesting a lower electron count consistent with the number of electrons in $Na^+$. Additionally, our choice of $Na^+$ was guided by the observation of heptacoordinated pentagonal bipyramidal geometry for one metal with the second metal having similar coordination environment, although based on the quality of electron density we modeled it in octahedral geometry. It is possible that the site is occupied by $Na^+$, by $Ca^{2+}$ with lower occupancy, or perhaps by a combination of both metals. 9PLC structure contains one DNA strand, one PEG modeled as two conformers, two potassium ions (one on a special position at 0.5 occupancy), two strontium ions and five water molecules. Despite high B-factors and a lack of visible coordinating partners, strontium ions were used in the final model due to significantly lower $R_{work}/R_{free}$ values (by ~1%) compared to when $Na^+$ or $K^+$ ions were placed in that density. Data collection and refinement statistics are presented in Table S3. Subsequent structural analysis utilized the algorithm ASC-G4[7] and 3DNA[8].

**Fluorescence anisotropy**

Pre-annealed FAM-labeled DNA was diluted to 10 nM in binding buffer containing 10 mM Tris-HCl, pH 7.4, and 100 mM KCl. Increasing concentrations of peptide were added to the DNA solution and incubated at 25 °C for 30–60 min in the dark before measurement. Fluorescence anisotropy was recorded using excitation at 492 nm and emission at 520 nm on a Cary Eclipse fluorescence spectrometer (Agilent Technologies, USA). The four polarization components, $I_{VV}$, $I_{VH}$, $I_{HV}$ and $I_{HH}$, were measured, and the G-factor was calculated as $G = I_{HV}/I_{HH}$. Fluorescence anisotropy was then calculated as $r = (I_{VV} – G × I_{VH}) / (I_{VV} + 2 × G × I_{VH})$. Binding curves were generated by plotting anisotropy values against peptide concentration and fitted with a one-site binding model to obtain apparent $K_d$ values. Assuming peptide binding and drLHG4 formation were coupled, the fitted values were interpreted as apparent folding-coupled affinities.

**Fluorescence titration of ZnTEG$^Q$PyrPc with DNA**

Fluorescence titrations of ZnTEG$^Q$PyrPc were performed by keeping the ligand concentration constant and titrating increasing concentrations of pre-annealed DNA in binding buffer containing 10 mM Tris-HCl, pH 7.4, and 100 mM KCl. All mixed samples were heated at 90 °C for 5 min and incubated at 25 °C for 30–60 min in the dark before measurement. Fluorescence spectra were recorded on a Cary Eclipse fluorescence spectrometer (Agilent Technologies, USA). Binding-response curves were generated by plotting the fluorescence change at the selected emission wavelength against total DNA concentration and fitted empirically with an apparent saturation equation, $\Delta F.I. = \Delta F.I._{max}[D]/(K_{1/2,app} + [D])$, where [D] is the total DNA concentration. Because ZnTEG$^Q$PyrPc is aggregated under the assay conditions and fluorescence enhancement reflects coupled aggregate disruption, monomer stabilization, and DNA binding/folding, $K_{1/2,app}$ was reported as an apparent half-maximal response concentration rather than a microscopic 1:1 dissociation constant.

**Table S1.** DNA sequences used in this study.

| Name | Sequence |
|---|---|
| $d(GT)_7$ | GTGTGTGTGTGTGT |
| $d(GT)_{7.5}$ | GTGTGTGTGTGTGTG |
| $d(GT)_8$ | GTGTGTGTGTGTGTGT |
| $d(GT)_9$ | GTGTGTGTGTGTGTGTGT |
| $d(GC)_9$ | GCGCGCGCGCGCGCGCGC |
| $d(GA)_6$ | GAGAGAGAGAGA |
| $d(GA)_7$ | GAGAGAGAGAGAGA |
| $d(GA)_{7.5}$ | GAGAGAGAGAGAGAG |
| $d(GA)_9$ | GAGAGAGAGAGAGAGAGA |
| $d(GA)_8$ | GAGAGAGAGAGAGAGA |
| $d(GA)_{10}$ | GAGAGAGAGAGAGAGAGAGA |
| $d(GA)_{11}$ | GAGAGAGAGAGAGAGAGAGAGA |
| $d(GT)_{16}$ | GTGTGTGTGTGTGTGTGTGTGTGTGTGTGTGT |
| $T_5$-$d(GT)_{7.5}$-$T_5$ | TTTTTGTGTGTGTGTGTGTGTTTTT |
| $T_5$-$d(GT)_{11.5}$-$T_5$ | TTTTTGTGTGTGTGTGTGTGTGTGTGTGTTTTT |
| $T_5$-$d(GT)_{15.5}$-$T_5$ | TTTTTGTGTGTGTGTGTGTGTGTGTGTGTGTGTGTGTTTTT |
| $d(GT)_5$ | GTGTGTGTGT |
| $d(GT)_6$ | GTGTGTGTGTGT |
| $d(GT)_{10}$ | GTGTGTGTGTGTGTGTGTGT |
| $d(GT)_{11}$ | GTGTGTGTGTGTGTGTGTGTGT |
| $d(GT)_{12}$ | GTGTGTGTGTGTGTGTGTGTGTGT |
| $d(GT)_{11.5}$ | GTGTGTGTGTGTGTGTGTGTGTG |
| $d(GT)_{15.5}$ | GTGTGTGTGTGTGTGTGTGTGTGTGTGTGTG |
| $d(GA)_{12}$ | GAGAGAGAGAGAGAGAGAGAGAGA |
| $d(GC)_{7.5}$ | GCGCGCGCGCGCGCG |
| TT-$d(GT)_{7.5}$-TT | TTGTGTGTGTGTGTGTGTT |

| Name | Sequence |
| --- | --- |
| 3Less246/d(GTG)$_4$/Motif1 | GTGGTGGTGGTG |
| 3Less135/Motif2 | GGTGGTGGTGTG |
| 1Less1 | GGTGTGTGTGTGTG |
| 1Less2 | GTGGTGTGTGTGTG |
| 1Less3 | GTGTGGTGTGTGTG |
| 1Less4 | GTGTGTGGTGTGTG |
| 1Less5 | GTGTGTGTGGTGTG |
| 1Less6 | GTGTGTGTGTGGTG |
| 1Less7 | GTGTGTGTGTGTGG |
| 1More1 | GTTGTGTGTGTGTGTG |
| 1More2 | GTGTTGTGTGTGTGTG |
| 1More3 | GTGTGTTGTGTGTGTG |
| 1More4 | GTGTGTGTTGTGTGTG |
| 1More5 | GTGTGTGTGTTGTGTG |
| 1More6 | GTGTGTGTGTGTTGTG |
| 1More7 | GTGTGTGTGTGTGTTG |
| 2More44 | GTGTGTGTTTGTGTGTG |
| 2More77 | GTGTGTGTGTGTGTTTG |
| 2Less13 | GGTGGTGTGTGTG |
| 2Less15 | GGTGTGTGGTGTG |
| 2Less17 | GGTGTGTGTGTGG |
| 2Less37 | GTGTGGTGTGTGG |
| 2Less57 | GTGTGTGTGGTGG |
| 2Less35 | GTGTGGTGGTGTG |
| 2Less24/Motif1-bulge1 | GTGGTGGTGTGTG |
| 2Less26/Motif1-bulge2 | GTGGTGTGTGGTG |
| 2Less46/Motif1-bulge3 | GTGTGTGGTGGTG |
| 2Less34 | GTGTGGGTGTGTG |

| Name | Sequence |
|---|---|
| 2Less45 | GTGTGTGGGTGTG |
| 2Less67 | GTGTGTGTGTGGG |
| 2Less27 | GTGGTGTGTGTGG |
| 2Less47 | GTGTGTGGTGTGG |
| 2Less25 | GTGGTGTGGTGTG |
| 2Less12 | GGGTGTGTGTGTG |
| 2Less14 | GGTGTGGTGTGTG |
| 2Less16 | GGTGTGTGTGGTG |
| 2Less36 | GTGTGGTGTGGTG |
| 3Less137 | GGTGGTGTGTGG |
| 3Less136 | GGTGGTGTGGTG |
| 3Less146 | GGTGTGGTGGTG |
| 3Less147 | GGTGTGGTGTGG |
| 3Less157 | GGTGTGTGGTGG |
| 3Less357 | GTGTGGTGGTGG |
| 3Less247 | GTGGTGGTGTGG |
| 3Less257 | GTGGTGTGGTGG |
| 3Less123 | GGGGTGTGTGTG |
| 3Less124 | GGGTGGTGTGTG |
| 3Less134 | GGTGGGTGTGTG |
| 3Less234 | GTGGGGTGTGTG |
| 3Less235 | GTGGGTGGTGTG |
| 3Less236 | GTGGGTGTGGTG |
| 3Less567 | GTGTGTGTGGGG |
| 2Less67G5 | GTGTGTGTGGTGGG |
| 2Less67G6 | GTGTGTGTGTGGGG |
| 2Less34G3 | GTGTGGGGTGTGTG |
| 2Less34G8 | GTGTGGGTGTGTGG |

| Name | Sequence |
| --- | --- |
| $d(GT)_{7.5}$-1toG | GGGTGTGTGTGTGTG |
| $d(GT)_{7.5}$-4toG | GTGTGTGGGTGTGTG |
| $d(GT)_{7.5}$-7toG | GTGTGTGTGTGTGGG |
| $d(GT)_{7.5}$-1toA | GAGTGTGTGTGTGTG |
| $d(GT)_{7.5}$-4toA | GTGTGTGAGTGTGTG |
| $d(GT)_{7.5}$-7toA | GTGTGTGTGTGTGAG |
| $d(GT)_{7.5}$-1toC | GCGTGTGTGTGTGTG |
| $d(GT)_{7.5}$-4toC | GTGTGTGCGTGTGTG |
| $d(GT)_{7.5}$-7toC | GTGTGTGTGTGTGCG |
| $d(GT)_{7.5}$-2C | GTGTGCGTGTGCGTG |
| $d(GT)_{7.5}$-3C | GTGCGTGCGTGCGTG |
| $d(GT)_{7.5}$-4C | GCGTGCGTGCGTGCG |
| $d(GA)_{7.5}$-2C | GAGAGCGAGAGCGAG |
| 4Less2467 | GTGGTGGTGGG |
| T-Motif1 | TGTGGTGGTGGTG |
| Motif1-4A | GAGGAGGAGGAG |
| Motif1-4C | GCGGCGGCGGCG |
| Motif1-T2A | GTGGAGGTGGTG |
| Motif1-T4A | GTGGTGGTGGAG |
| Motif1-T2TT | GTGGTTGGTGGTG |
| Motif1-T4TT | GTGGTGGTGGTTG |
| Yeast14 | GGTGTGTGGGTGTG |
| NPM-C70_F | GATC**GAATTC**CAAGAATCCTTC |
| NPM-C70_R | CATG**AAGCTT**TTAAAGAGACTT |

**Table S2.** Thermal stability of triple-deletion d$(GT)_{7.5}$-derived variants.

| Name | Sequence | $T_m$ (°C) |
|---|---|---|
| 3Less246 | GTGGTGGTGGTG | >90 |
| 3Less135 | GGTGGTGGTGTG | 73.5 |
| 3Less137 | GGTGGTGTGTGG | >90 |
| 3Less136 | GGTGGTGTGGTG | 37.3 |
| 3Less146 | GGTGTGGTGGTG | 38.9 |
| 3Less147 | GGTGTGGTGTGG | 34.8 |
| 3Less157 | GGTGTGTGGTGG | >90 |
| 3Less357 | GTGTGGTGGTGG | 76.4 |
| 3Less247 | GTGGTGGTGTGG | / |
| 3Less257 | GTGGTGTGGTGG | / |
| 3Less123 | GGGGTGTGTGTG | / |
| 3Less124 | GGGTGGTGTGTG | 48.7 |
| 3Less134 | GGTGGGTGTGTG | 54.5 |
| 3Less234 | GTGGGGTGTGTG | / |
| 3Less235 | GTGGGTGGTGTG | 63.5 |
| 3Less236 | GTGGGTGTGGTG | 55.2 |
| 3Less567 | GTGTGTGTGGGG | / |

/ = not folded or $T_m < 30$ °C

**Table S3.** Crystal data and structure refinement for *2Less26* crystals

| | **$SrCl_2$** | **$CaCl_2$** |
|---|---|---|
| **Resolution range, Å** | 57.044–1.538 | 23.089–0.878 |
| **Highest resolution shell, Å** | 1.565–1.538 | 0.893-0.878 |
| **Space group** | $P6_522$ | $P12_11$ |
| **Unit cell dimensions** | | |
| ***a*, *b*, *c* (Å)** | 30.90, 30.90, 114.90 | 24.6, 46.2, 28.3 |
| ***α*, *β*, *γ* (°)** | 90, 90, 120 | 90, 115.6, 90 |
| **Unique reflections** | 5136 | 2241 |
| **Redundancy** | 34.4 (37.5) | 4.0 (4.0) |
| **Completeness (%)** | 95.8 (100) | 100 (100) |
| ***I*/sigma** | 18.9 (1.3) | 7.6 (1.4) |
| ***R*-merge** | 0.074 (2.965) | 0.104 (1.280) |
| **$R_{work}$/$R_{free}$ (%)** | 16.30/19.35 | 14.60/15.60 |
| **Molecular entities in ASU** | 1 | 2 |
| **Number of atoms** | 481 | 1016 |
| **DNA**<br>**Water**<br>**Potassium ion**<br>**Sodium ion**<br>**Strontium ion**<br>**PEG** | 424<br>5<br>2<br>0<br>2<br>48* | 845<br>132<br>3<br>2<br>0<br>31 |
| **Overall B-factor for ASU (Å$^2$)** | 51.79 | 9.46 |
| **RMS deviations** | | |
| **Bond length (Å)** | 0.012 | 0.010 |
| **Bond angles (°)** | 1.183 | 1.188 |
| **PDB ID** | **9PLC** | **9PLB** |

*This PEG molecule is present in two conformations and is shorter than that in the 9PLB structure.

**Table S4.** Out-of-plane deviations ($D_{OOP}$) in Å. Tetrad numbering is shown in the schematics below. Chain A is colored cyan and chain B is colored pink.

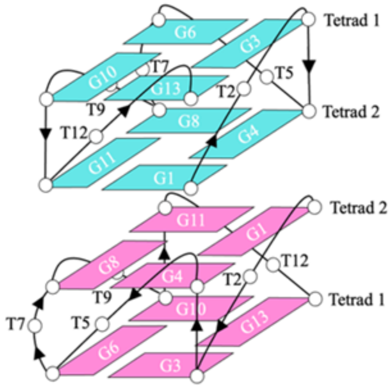


| | $D_{OOP}$ (Å) | |
|---|---|---|
| **Tetrad** | **9PLB** | **9PLC*** |
| **Chain A, Tetrad 1** | 1.23 | 1.12 |
| **Chain A, Tetrad 2** | 1.07 | 1.27 |
| **Chain B, Tetrad 1** | 1.34 | - |
| **Chain B, Tetrad 2** | 1.11 | - |

*In 9PLC Tetrad 1 is defined as having a thymine cap and Tetrad 2 is defined as the dimer interface (matching the definition for the 9PLB structure).

**Table S5.** Average groove width in the two *2Less26* structures measured from C5′-C5′ and reported in Å. The definition of chains and tetrads is the same as in Table S4.

**Table S5A.** Average groove widths for molecules A and B in 9PLB crystal structure.

| Groove between strands #-# | | **1-2** | **2-3** | **3-4** | **4-1** |
|---|---|---|---|---|---|
| **Tetrads (Å)** | Chain A, Tetrad 1 | 15.52 | 14.08 | 14.89 | 14.34 |
| | Chain A, Tetrad 2 | 14.84 | 15.64 | 15.14 | 15.69 |
| | Chain B, Tetrad 1 | 15.11 | 15.66 | 15.08 | 15.83 |
| | Chain B, Tetrad 2 | 15.27 | 14.10 | 15.27 | 14.15 |
| **Average (Å)** | | **15.2 ± 0.2** | **14.9 ± 0.8** | **15.1 ± 0.1** | **15.0 ± 0.8** |

**Table S5B.** Average groove widths for 9PLC.

| Groove between strands #-# | | **1-2** | **2-3** | **3-4** | **4-1** |
|---|---|---|---|---|---|
| **Tetrads (Å)** | Chain A, Tetrad 1 | 15.56 | 13.78 | 15.00 | 14.36 |
| | Chain A, Tetrad 2 | 15.23 | 15.18 | 15.22 | 15.58 |
| **Average (Å)** | | **15.4 ± 0.2** | **14.5 ± 0.7** | **15.1 ± 0.1** | **15.0 ± 0.6** |

**Table S6.** Helical angles (in degrees, °) between subsequent guanines in adjacent tetrads, per chain and strand.

| **Between G-tetrads:** | **Chain, Strand** | **9PLB** | **9PLC** |
|---|---|---|---|
| **Chain A, 1 and 2** | 1-2 | -26.2 | -26.0 |
| | 2-3 | -25.3 | -25.8 |
| | 3-4 | -26.5 | -25.6 |
| | 4-1 | -25.8 | -26.0 |
| Average | | **-26.0 ± 0.5** | **-25.9 ± 0.2** |
| **Chain A - Chain B** | 1-2 | -11.5 | |
| | 2-3 | -10.9 | |
| | 3-4 | -11.1 | |
| | 4-1 | -10.5 | |
| Average | | **-11.0 ± 0.4** | |
| **Chain B, 1 and 2** | 1-2 | -23.4 | |
| | 2-3 | -25.6 | |
| | 3-4 | -23.6 | |
| | 4-1 | -25.5 | |
| Average | | **-24.5 ± 1.0** | |

**Table S7.** G4 binding peptides used in this study.

| Name | Motif (N→C) |
|---|---|
| RHAU23 (23 aa) | HPGHLKGREIGMWYAKKQGQKNK |
| WRN-RQC (24 aa) | HSLFGTGKDQTESWWKAFSRQLIT |
| KR12C (12 aa) | KRIVKLIKKWLR |
| CIRBP-peptide12 (25 aa) | RGGSAGGRGFFRGGRGRGRGFSRGG |
| G4P (64 aa + His Tag) | HPGHLKGREIGMWYAKKQGQKNKGTGSGAGTGSGAGTGSGAHPGHLKGREIGMWYAKKQGQKNKHHHHHH |
| NPM-C70 (70 aa + His Tag) | HHHHHHQESFKKQEKTPKTPKGPSSVEDIKAKMQASIEKGGSLPKVEAKFINYVKNCFRMTDQEAIQDLWQWRKSL |

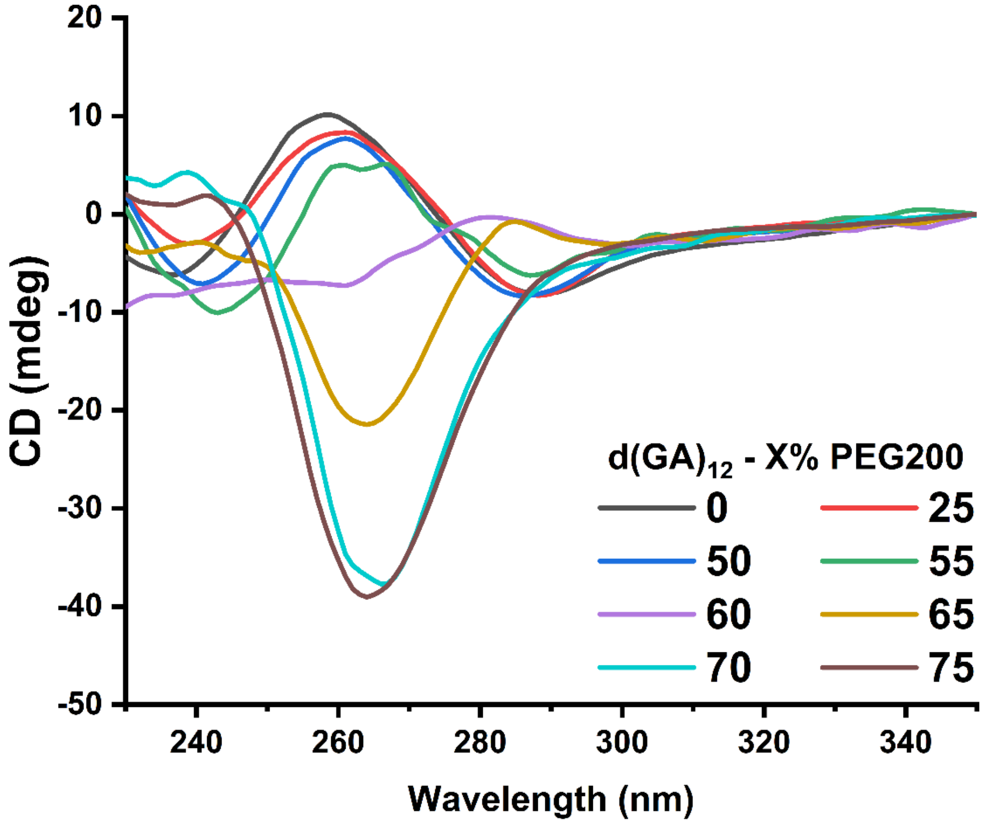


**Figure S1.** CD spectra of 5 μM d(GA)$_{12}$ measured at increasing PEG200 concentrations, showing progressive induction of the characteristic LHG4 signature.

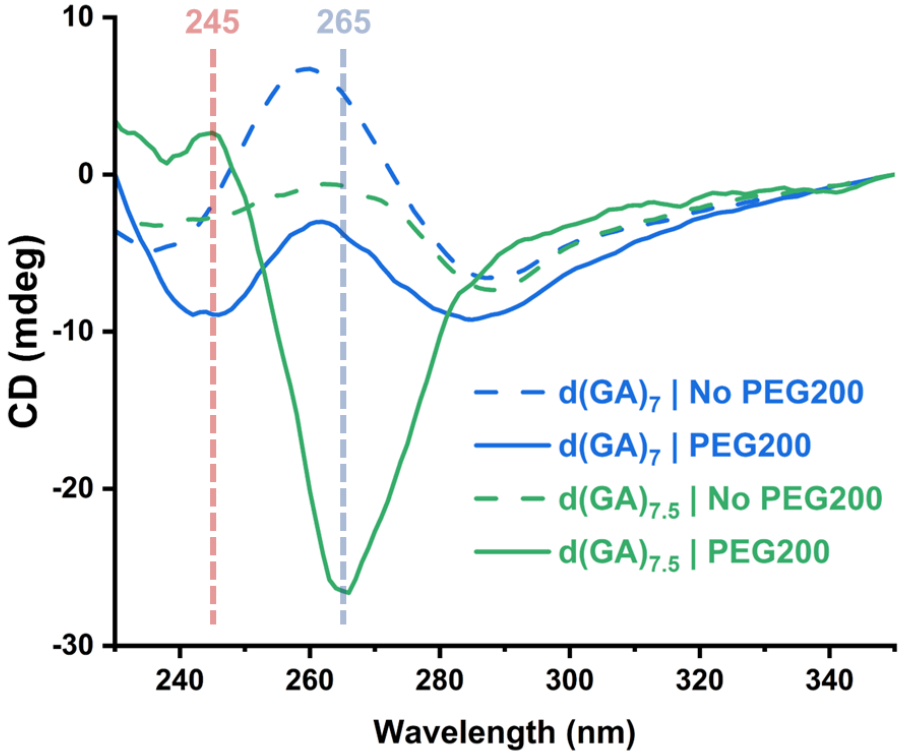


**Figure S2.** CD spectra of 5 μM d(GA)$_7$ and d(GA)$_{7.5}$ in the absence and presence of 75% PEG200 (v/v). Dashed lines indicate spectra recorded without PEG200, and solid lines indicate spectra recorded in PEG200.

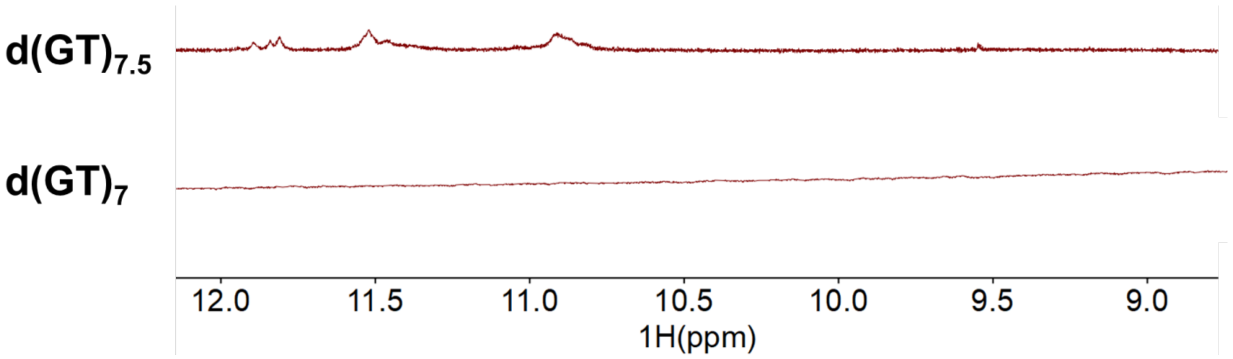


**Figure S3.** $^{1}$H NMR spectra of imino (12-9 ppm) peaks of 200 μM d(GT)$_{7.5}$ and d(GT)$_{7}$ with 50% DMSO.

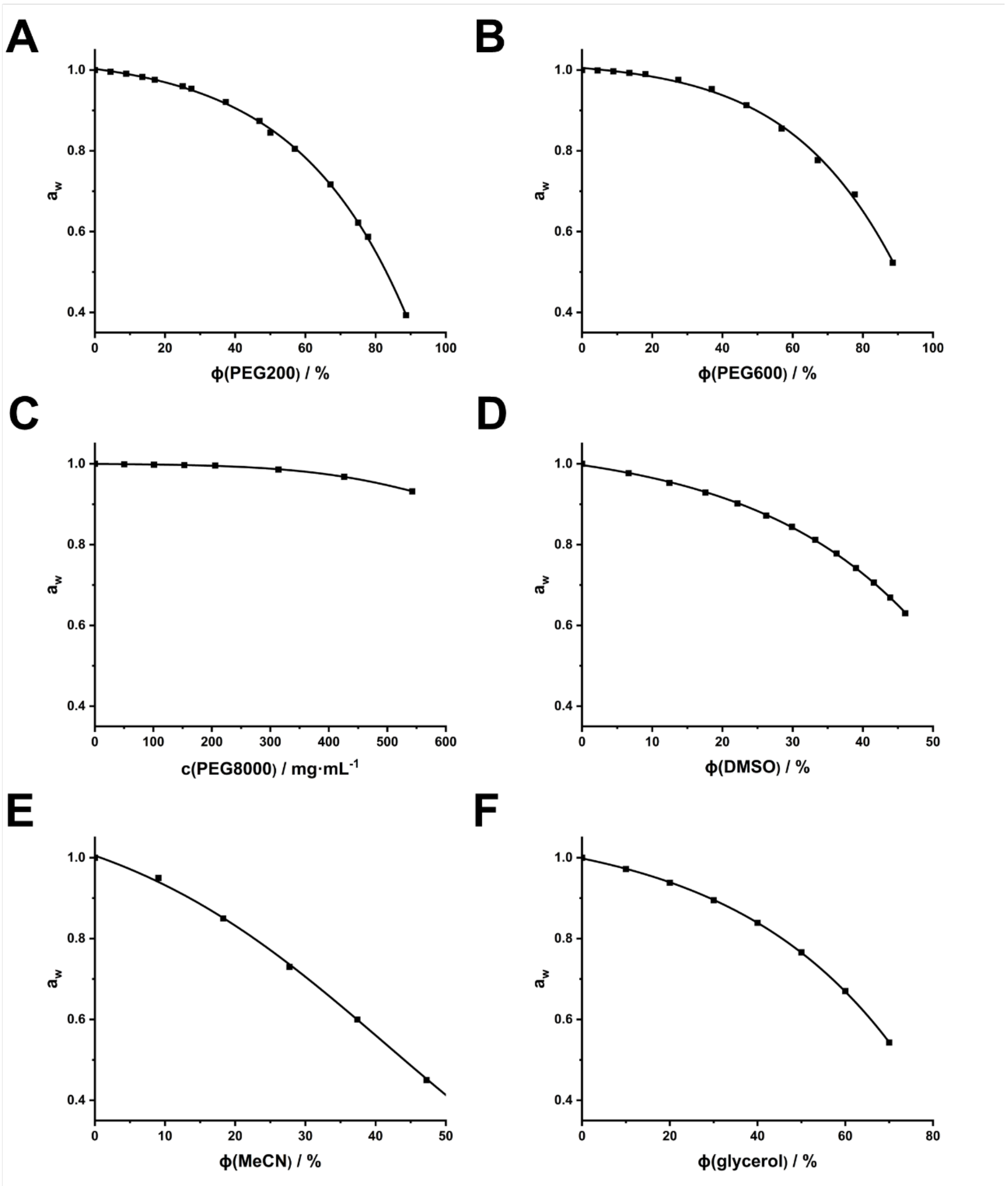


**Figure S4.** Water activity of cosolvent and crowding-agent solutions. Water activity ($a_w$) was measured at 25 °C using an AQUALAB TDL 2 water activity meter equipped with a tunable diode laser sensor. Values are plotted as a function of solute concentration for PEG200 (A), PEG600 (B), PEG8000 (C), DMSO (D), acetonitrile (MeCN) (E) and glycerol (F). PEG200, PEG600, DMSO, MeCN, and glycerol concentrations are expressed as % (v/v), whereas PEG8000 concentration is expressed as mg/mL. Water activity ($a_w$) is a dimensionless quantity expressed as the ratio of the equilibrium water vapor pressure above a solution to that above pure water at the same temperature, with pure water assigned an $a_w$ of 1.

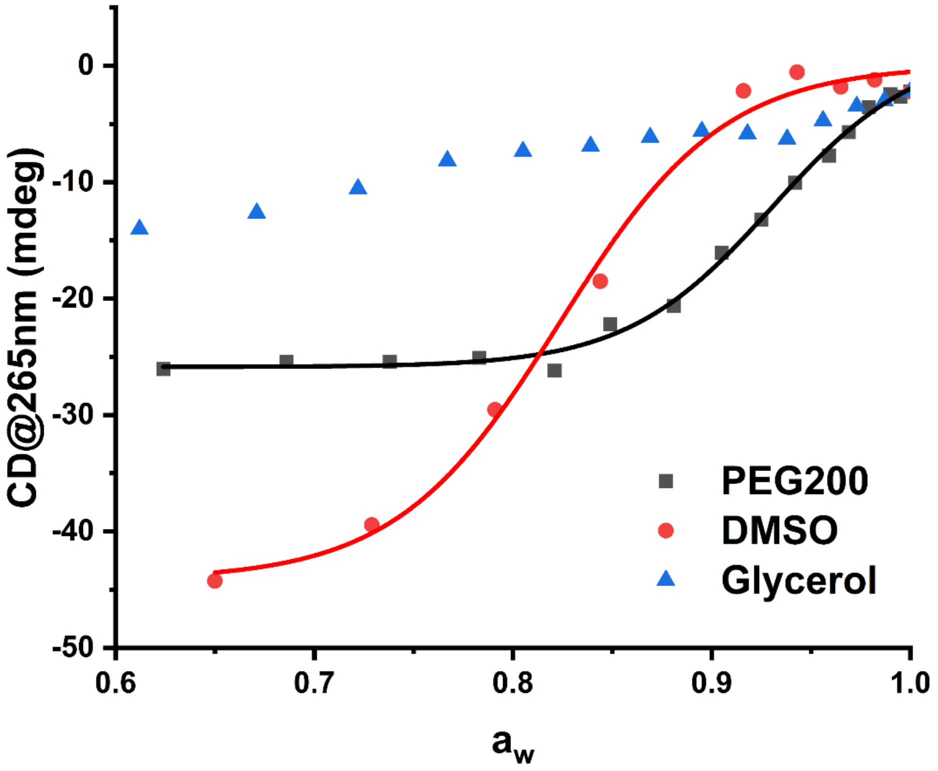


**Figure S5.** Relationship between bulk water activity and d(GT)$_{7.5}$ LHG4 induction. CD intensity at 265 nm for d(GT)$_{7.5}$ plotted as a function of water activity ($a_w$) in PEG200, DMSO and glycerol solutions. PEG200 and DMSO show sigmoidal transitions consistent with dehydration-induced LHG4 folding, whereas glycerol does not produce a comparable transition despite reducing bulk water activity.

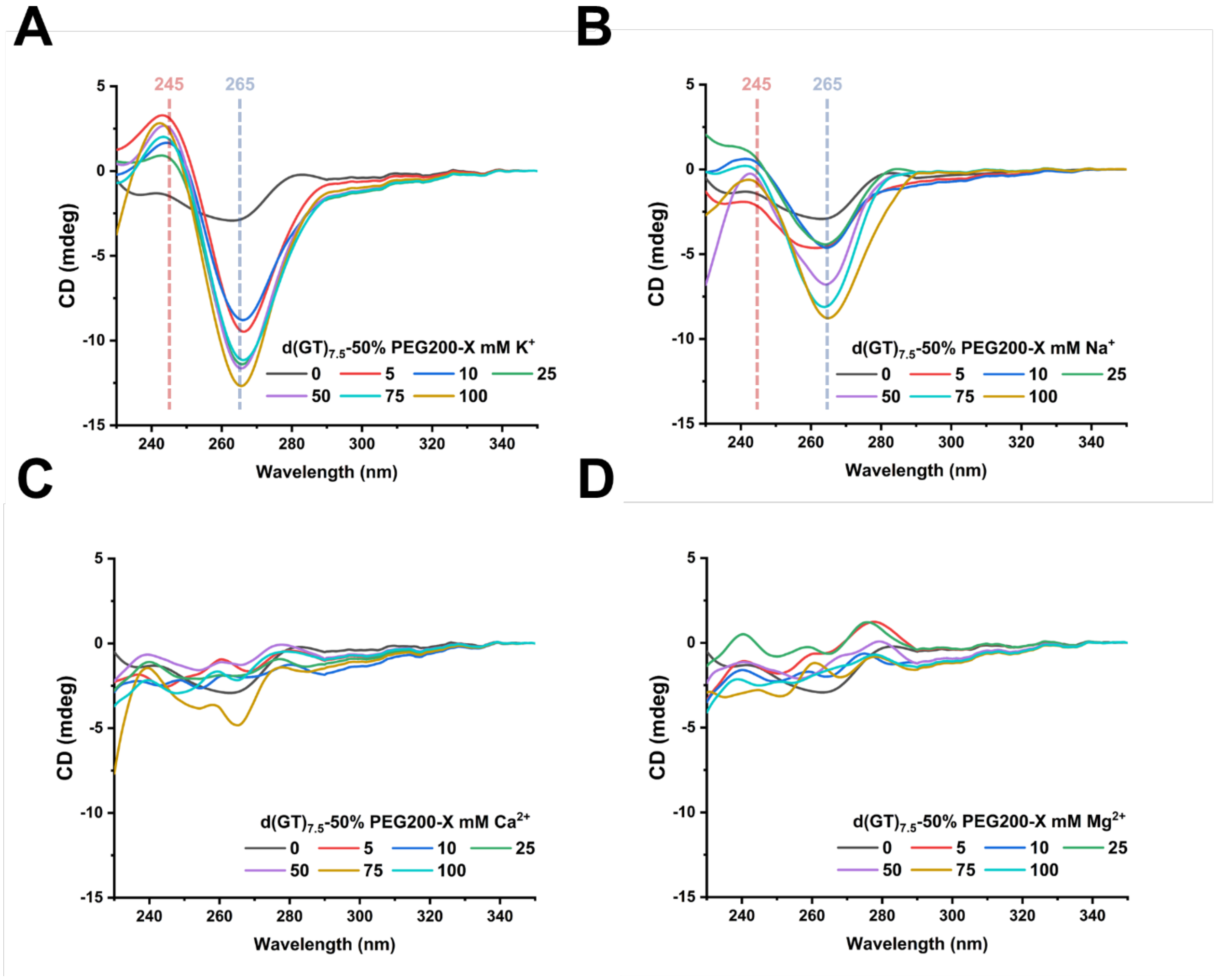


**Figure S6.** Ion dependence of d(GT)$_{7.5}$ LHG4 folding under dehydrating conditions. CD spectra of 5 μM d(GT)$_{7.5}$ measured in 50% PEG200 with increasing concentrations of $K^+$ (A), $Na^+$ (B), $Ca^{2+}$ (C) or $Mg^{2+}$ (D).

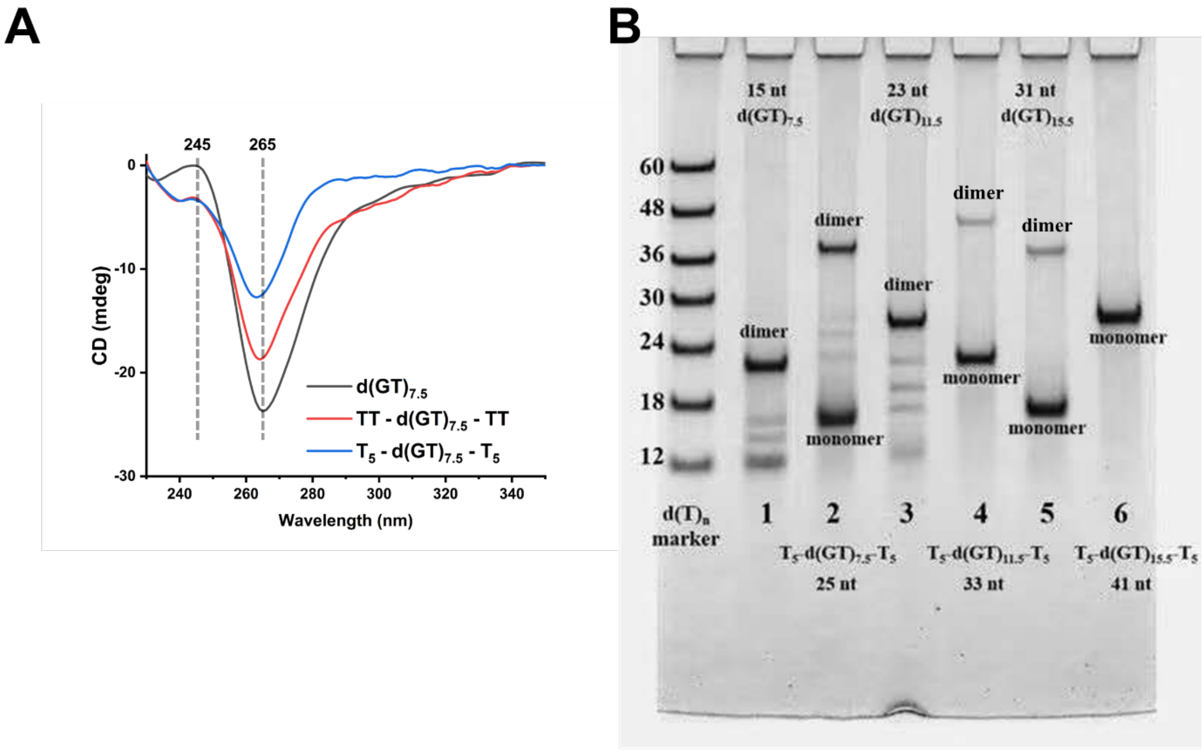


**Figure S7.** (A) Effect of terminal flanking sequences on $d(GT)_{7.5}$ LHG4 folding. CD spectra of $d(GT)_{7.5}$, TT-$d(GT)_{7.5}$-TT and $T_5$-$d(GT)_{7.5}$-$T_5$ measured under dehydrating conditions. Addition of terminal thymine flanks reduces the intensity of the characteristic LHG4 CD signal, with the strongest attenuation observed for the $T_5$-flanked sequence. Dashed vertical lines indicate 245 and 265 nm. (B) Native PAGE analysis of $d(GT)_n$ constructs with and without T5 flanking sequences. DNA samples were pre-folded at 5 μM in 50% PEG200 before loading, and the loading buffer also contained 50% PEG200 to minimize disruption of folded species during electrophoresis. Slow-migrating bands are consistent with dimeric or stacked two-block species, whereas faster-migrating bands correspond to monomeric species.

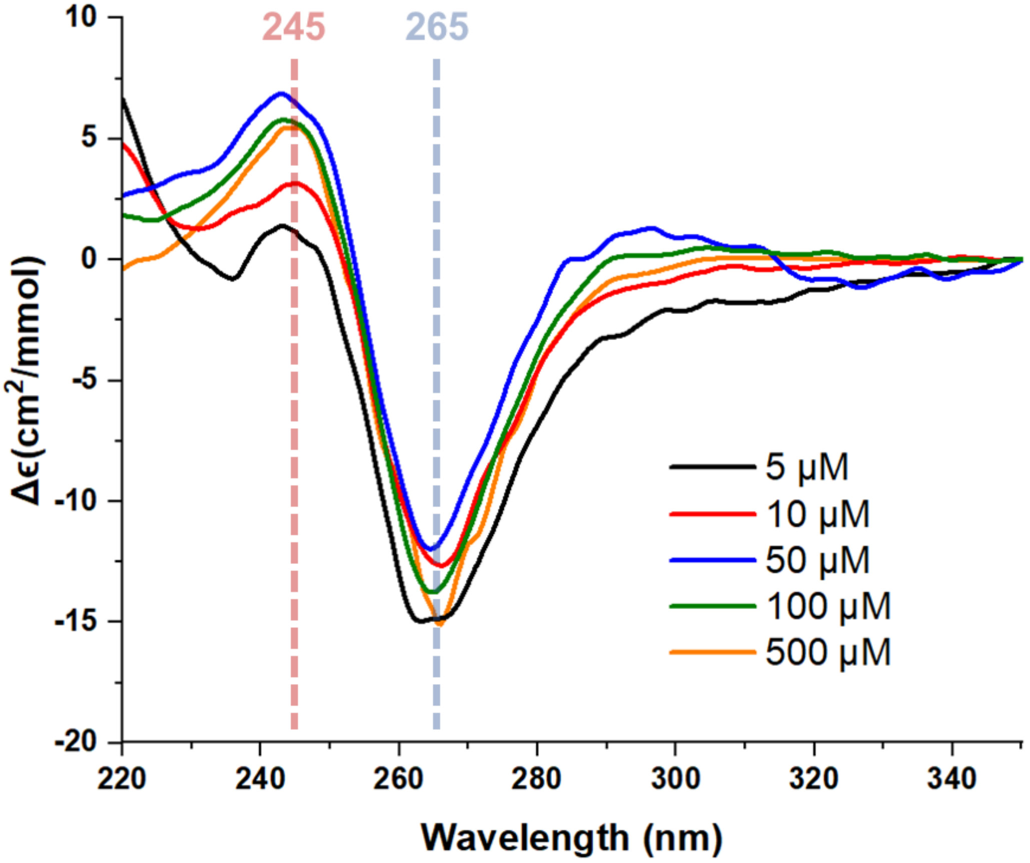


**Figure S8.** Concentration dependence of $d(GT)_{7.5}$ LHG4 folding. Molar CD spectra of $d(GT)_{7.5}$ measured at increasing DNA concentrations under 75% PEG200. The spectra show broadly similar LHG4 features across the tested concentration range, including a positive band near 245 nm and a negative band near 265 nm. The limited concentration dependence indicates that LHG4 folding is readily achieved under these conditions and is not strongly dependent on micromolar DNA concentration. Spectra at 5, 10, and 50 μM were recorded using a 10 mm quartz cuvette, whereas spectra at 100 and 500 μM were recorded using a 1 mm quartz cuvette.

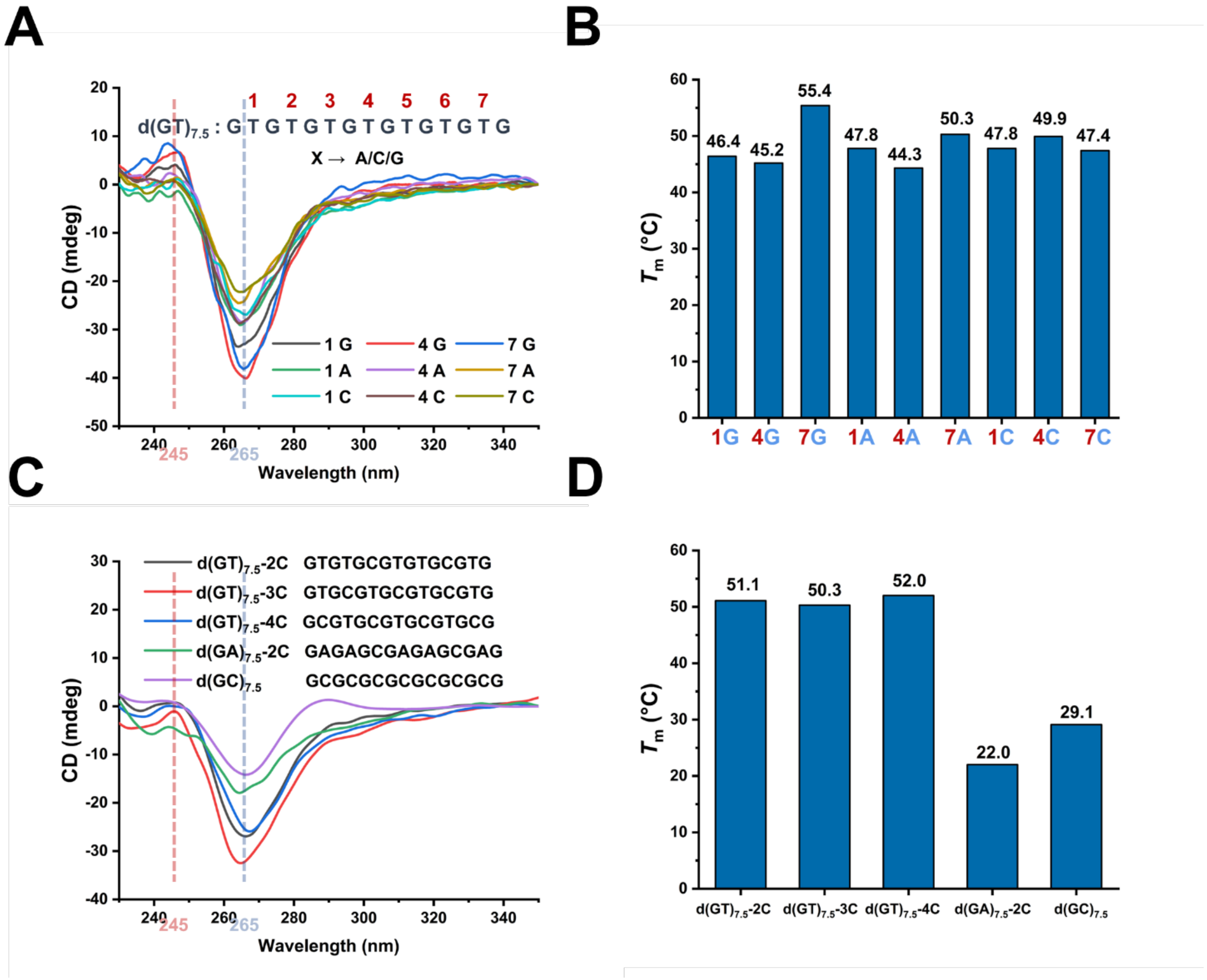


**Figure S9.** Effects of loop-base substitution on $d(GT)_{7.5}$-derived LHG4 folding. (A) CD spectra of 5 μM $d(GT)_{7.5}$ variants in which selected T residues were substituted with A, C or G. The position of the substituted base is indicated above the sequence. Most substitutions retained the characteristic LHG4-like CD signature, with a positive band near 245 nm and a negative band near 265 nm. (B) Melting temperatures of the corresponding single-substitution variants. (C) CD spectra of 5 μM variants containing multiple C substitutions, together with $d(GA)_{7.5}$-2C and $d(GC)_{7.5}$. (D) Melting temperatures of the corresponding variants.

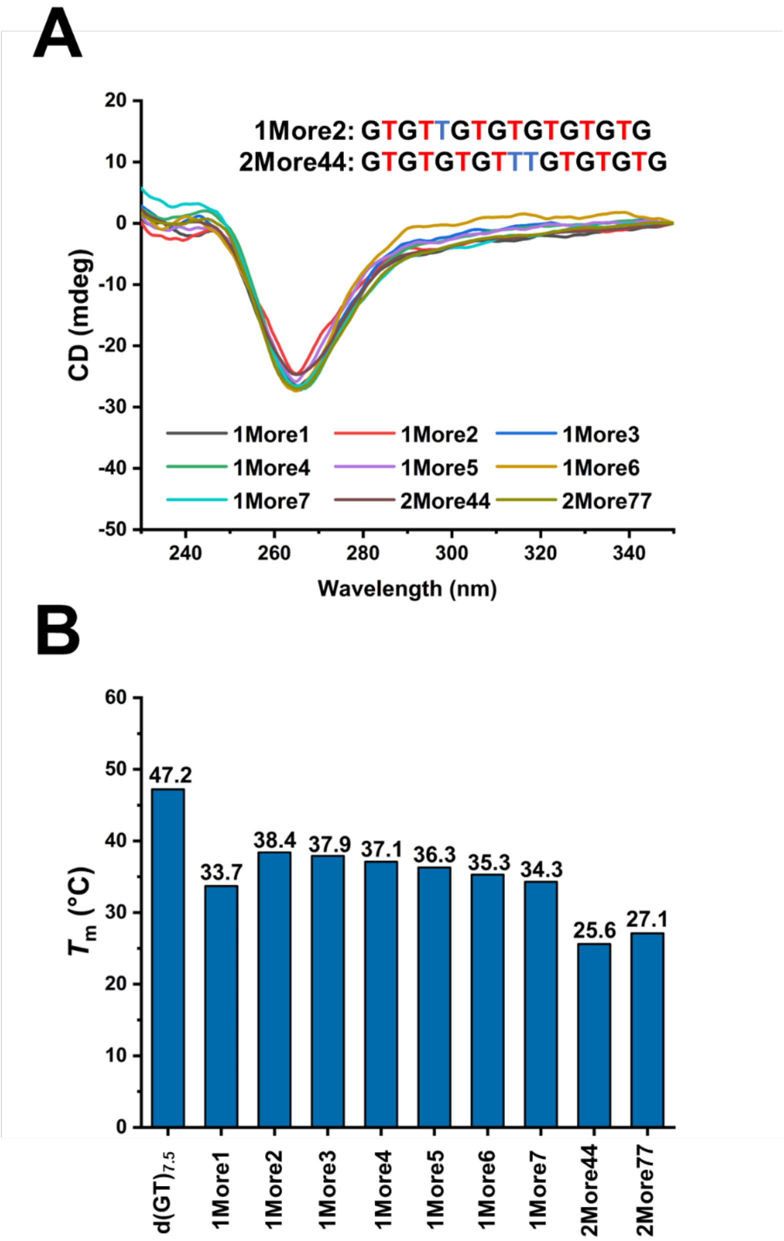


**Figure S10.** Effects of single- and double-nucleotide insertions on d(GT)$_{7.5}$-derived LHG4 folding. (A) CD spectra of 5 μM *1More* and *2More* variants under 75% PEG200 (25 °C), in which one or two additional T residues were introduced into the d(GT)$_{7.5}$ scaffold. All tested variants retained an LHG4-like CD profile, but with reduced thermal stability compared with d(GT)$_{7.5}$. (B) Melting temperatures of d(GT)$_{7.5}$, *1More* variants and *2More* variants. The inserted residues generally destabilized the fold, with the strongest effects observed for *2More44* and *2More77*.

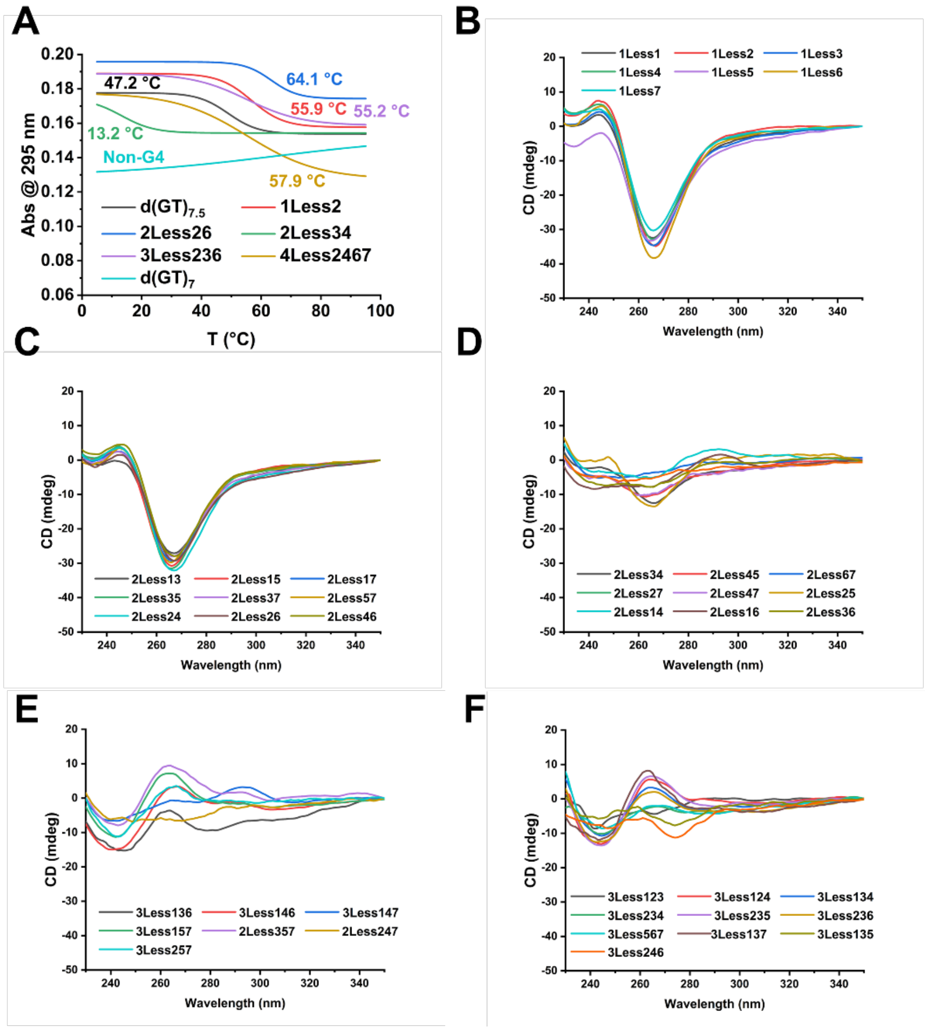


**Figure S11.** Deletion-dependent remodeling of d(GT)$_{7.5}$-derived LHG4 folding. (A) UV melting profiles of d(GT)$_{7.5}$ and selected deletion variants at 5 μM, with apparent $T_m$ values indicated on the curves. Selected deletions increased thermal stability, whereas others strongly destabilized the folded state or did not yield a G4-like transition. (B) CD spectra of 5 μM single-deletion variants (*1Less1*–*1Less7*) under 75% PEG200 (25 °C), showing that removal of one T residue is generally compatible with LHG4-like folding. (C) CD spectra of even-pattern double-deletion variants, which retain strong LHG4-like CD signatures. (D) CD spectra of odd-pattern double-deletion variants, which show weaker or noncanonical CD profiles. (E, F) CD spectra of representative triple-deletion variants.

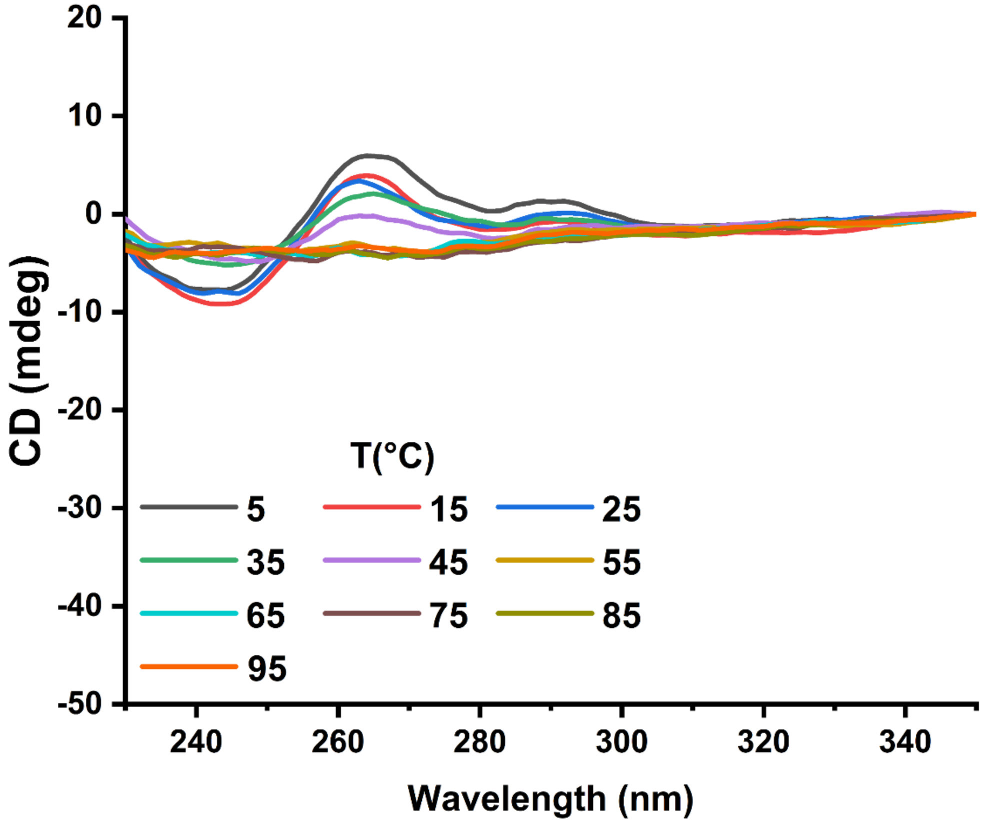


**Figure S12.** Temperature-dependent CD melting of the CD-weak *3Less236*. CD spectra recorded at increasing temperatures from 5 to 95 °C under 75% PEG200.

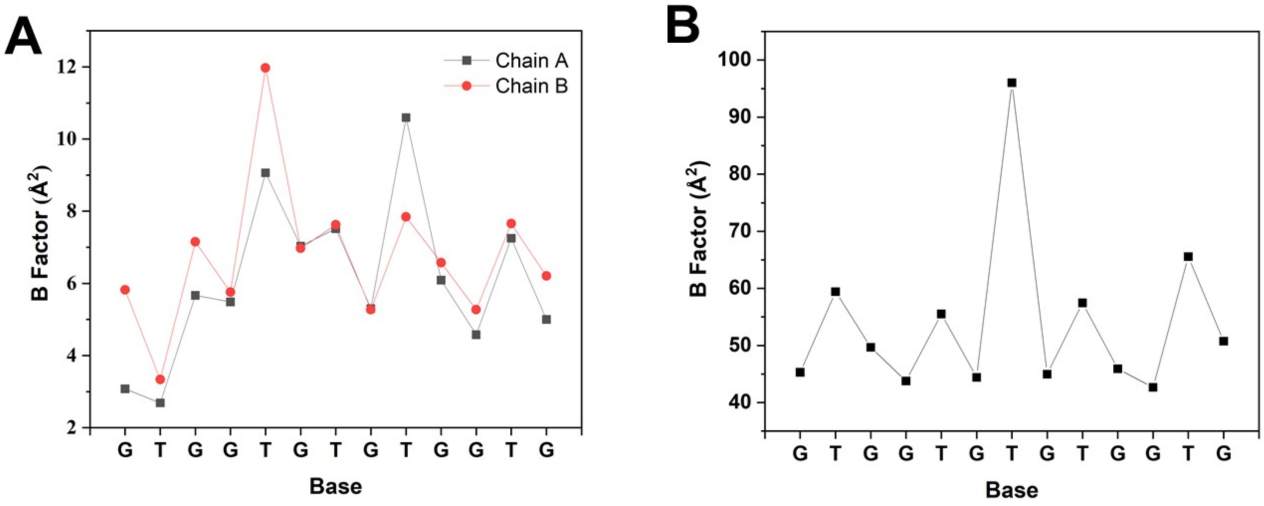


**Figure S13.** Comparison of B-factors by nucleotide for (**A**) 9PLB and (**B**) 9PLC structures. The significant differences in B-factors are due to differences in the resolution: 0.88 Å for 9PLB and 1.54 Å for 9PLC. The highest B values are observed for nucleotides facing the solvent: T9 of 9PLB Chain A, T5 of 9PLB Chain B, and T7 of 9PLC.

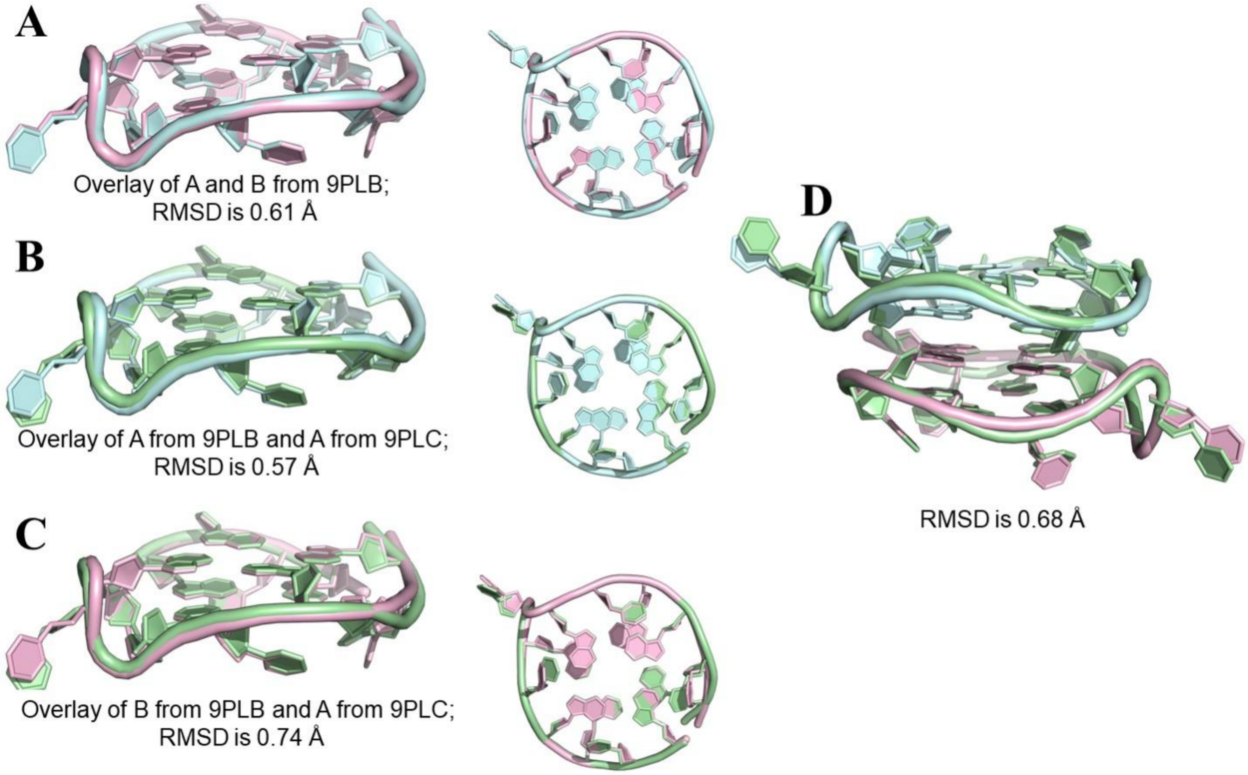


**Figure S14.** Overlays of *2Less26* structures. (**A**) Overlay of Chain A (blue) and B of 9PLB (light blue). Side (left) and Top (right) view. (**B**) Overlay of Chain A from 9PLB (blue) with Chain A of 9PLC (pink). (**C**) Overlay of Chain B from 9PLB (blue) with Chain A of 9PLC (pink). (D) Overlay of 9PLB (blue and pink) with the symmetry generated dimer of 9PLC (green).

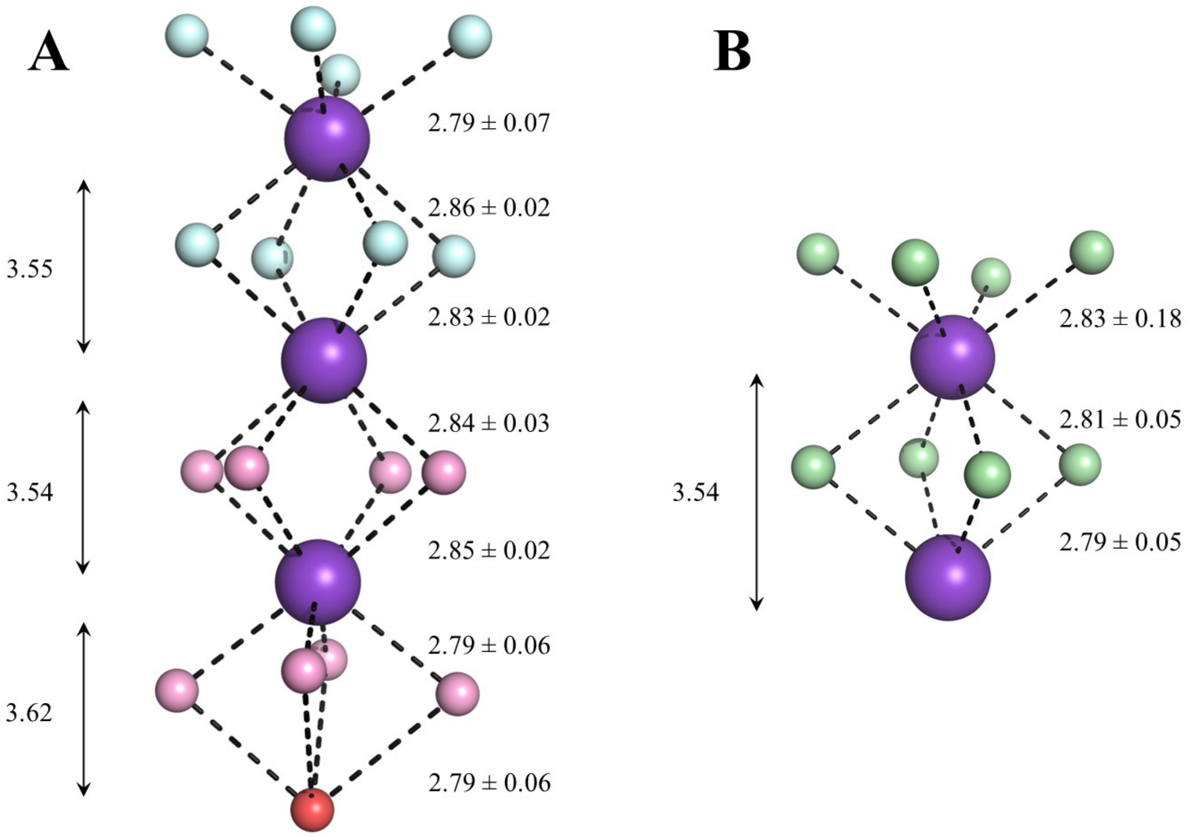


**Figure S15.** Ion channel in (**A**) 9PLB and (**B**) 9PLC structures. Cyan oxygen atoms correspond to molecule A and salmon oxygen atoms correspond to molecule B in 9PLB, while green oxygen atoms correspond to molecule A in 9PLC. Potassium ions are shown as purple spheres. Water molecule in 9PLB is colored red. Estimated standard deviations are given for the four averaged K-O distances. K2 in both structures is at the dimer interface. Distances are given in Å.

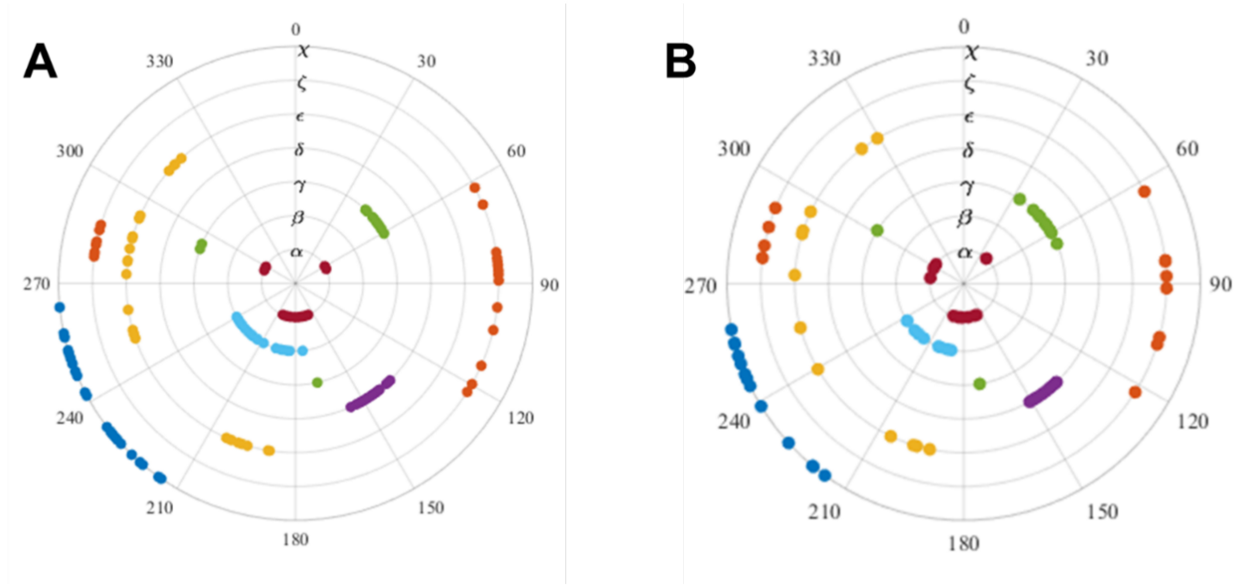


**Figure S16.** Torsion angle (°) wheel plots for (**A**) 9PLB and (**B**) 9PLC. Values are given only for the major conformation.

### Intermolecular contacts in *2Less26* crystals

In both 2*Less*26 structures, the crystal lattice is formed by contacts between different DNA chains via their thymine loops or bulges, cations and PEG200. In subsequent descriptions the symmetry-generated molecule is denoted by ', with multiple 's used to distinguish different symmetry-generated chains. In 9PLB, T7 of Chain A, T7' and T2'' of Chain B form a three-layered stack (Figure S17A); T5 of Chain A and T9''' of Chain B form an offset T-T base pair with two reciprocal hydrogen bonds between O2 and N3 (Figure S17B). In 9PLC, T2 and T9' (as well as T2' and T9) and T7 and T7'' stack (Figure S17C). Through these thymines, each 9PLB and 9PLC dimer contacts five and six symmetry molecules, respectively.

Metal ions and PEG200 were present in both structures, with more extensive contacts in 9PLB, consistent with its higher resolution. In 9PLB, two sodium ions link the phosphate backbones of neighboring dimers and adopt hexa- or heptacoordinated geometries completed by structural water molecules (Figure S17D). In 9PLC, two $Sr^{2+}$ ions occupy the space between DNA molecules and one coordinates N3 of T5 at 3.39 Å. These ions are likely hydrated, although their hydration shells are not resolved.

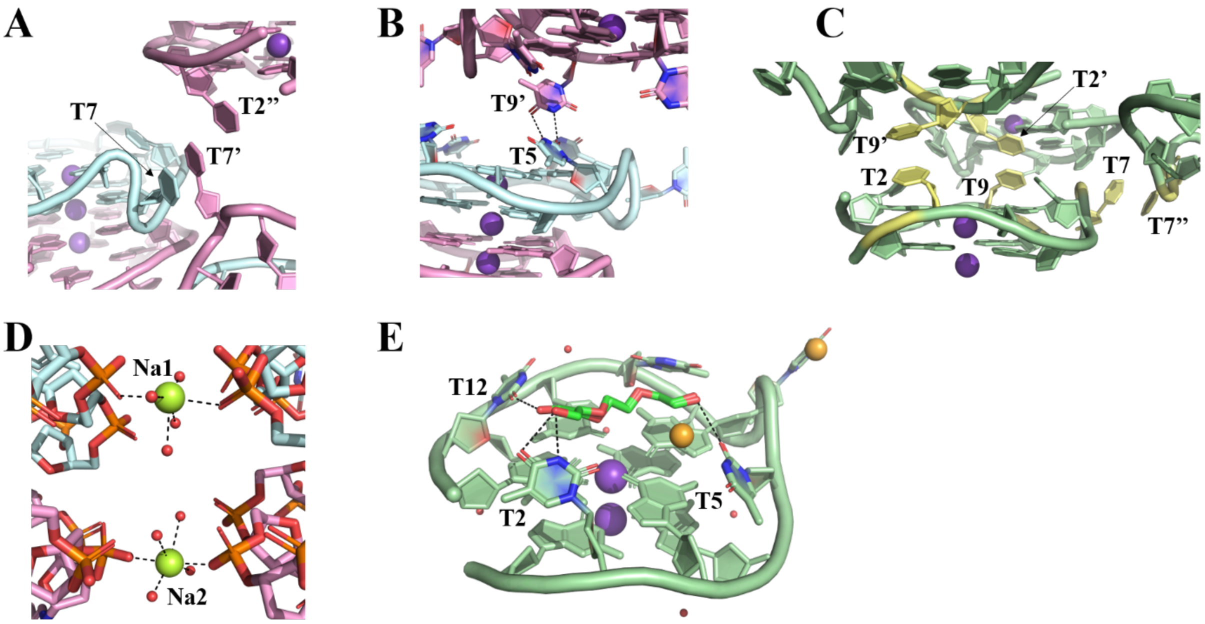

**Figure S17**. **3D organization of DNA, ions, and PEG molecules in *2Less26* crystals**. (**A**) Three-layered thymine stack (T7, T7' and T2'') in 9PLB structure. Two ASUs are shown. (**B**) T5-T9' base pair in 9PLB structure. Thymine oxygens and nitrogens are colored red and blue, respectively. (**C**) Stacking interactions in 9PLC between thymines shown in yellow for clarity. (**D**) Coordination of sodium ions in 9PLB structure. Waters are shown in red and $Na^{+}$ ions in green. (**E**) Interactions involving PEG200 molecule in 9PLC structure. PEG is colored by atom: carbons are forest green while oxygens are red. Orange spheres are $Sr^{2+}$ ions.

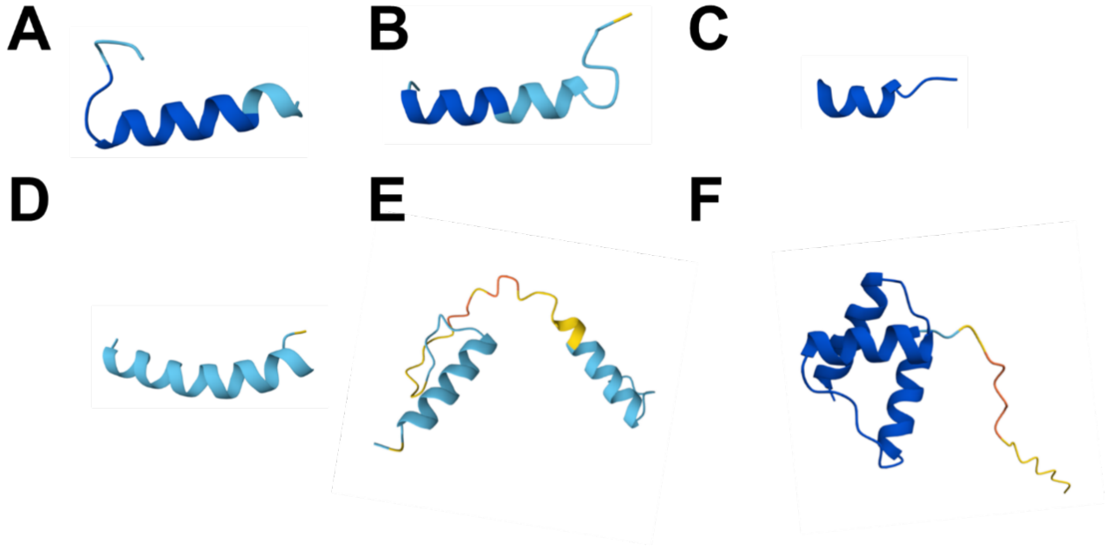


**Figure S18.** Predicted structural models of the protein-derived G4-binding peptides and domains screened in this study. Models were generated using AlphaFold 3 via AlphaFold Server. (A) RHAU23; (B) WRN-RQC; (C) KR12C; (D) CIRBP-peptide12; (E) G4P; and (F) NPM-C70. Amino acid sequences are provided in Table S7.

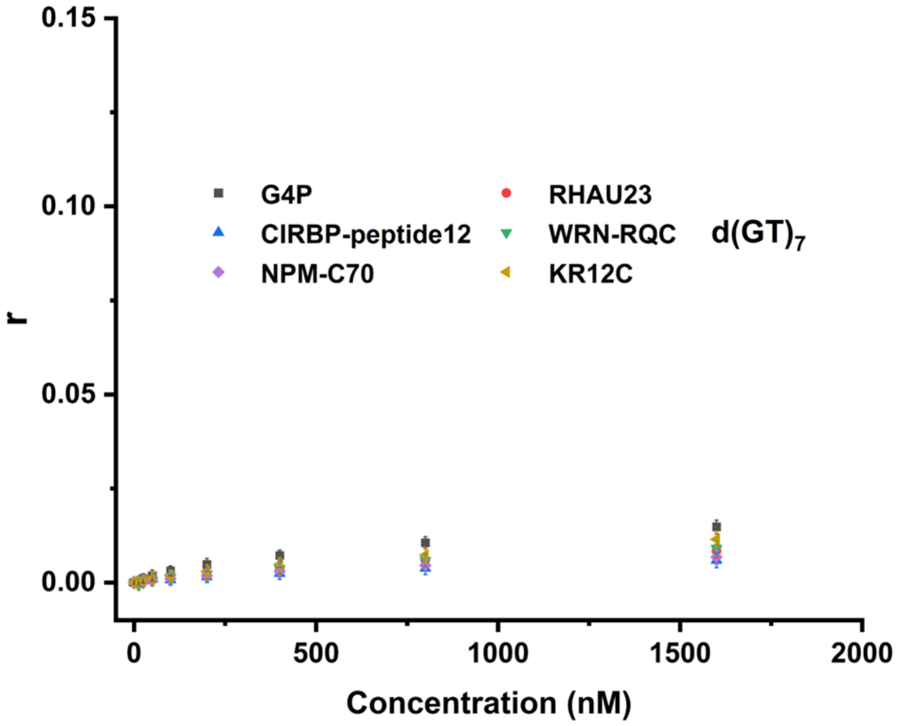


**Figure S19.** Fluorescence anisotropy analysis of interactions between protein-derived G4-binding fragments/domains and $d(GT)_7$ DNA. None of the tested binders produced a reliable concentration-dependent increase in anisotropy under these conditions, and no apparent $K_d$ could be determined.

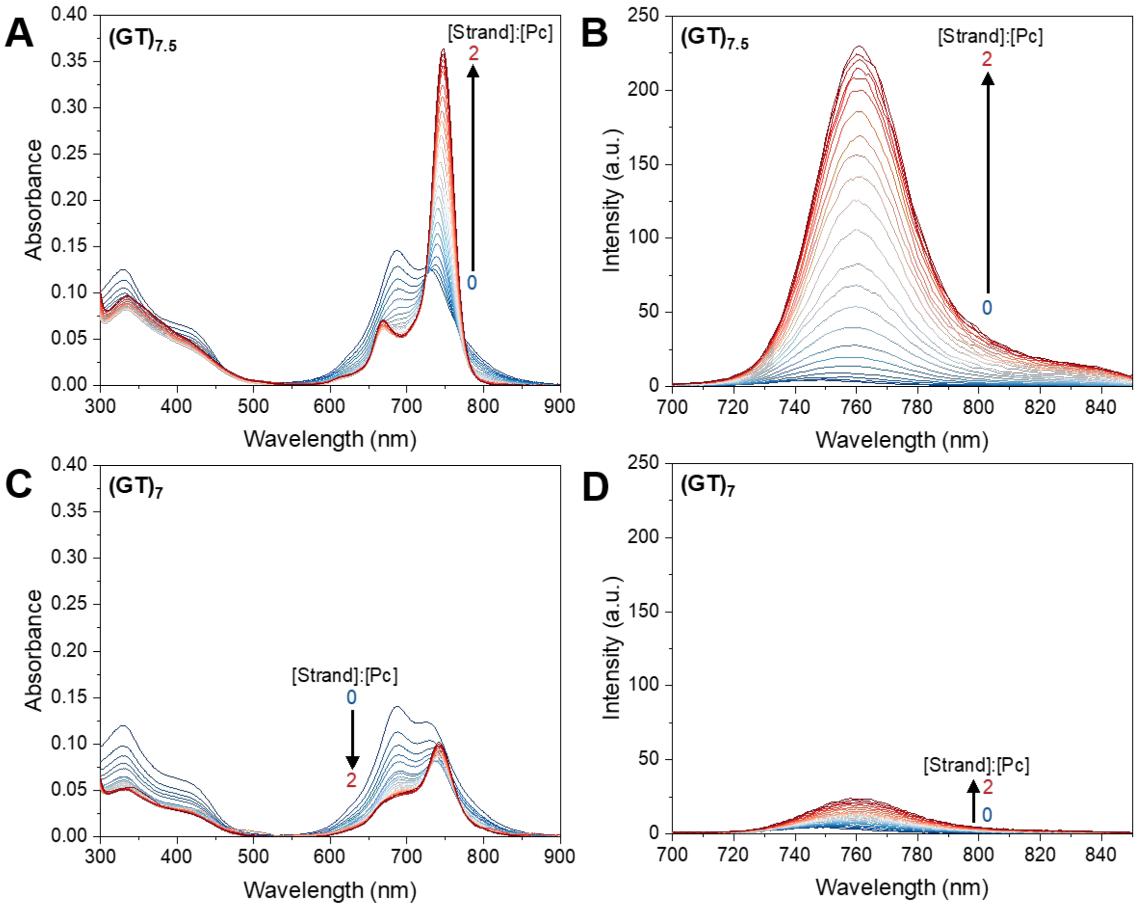


**Figure S20.** UV-visible absorption and emission spectra of ZnTEG$^{Q}$PyrPc (2.3 μM) in buffer (30 mM Tris-HCl, pH 7.4, 50 mM $K^+$) with increasing concentration of (A, B) d(GT)$_{7.5}$ and (C, D) d(GT)$_7$ (0-4.3 μM). These data indicate that d(GT)$_{7.5}$ promotes phthalocyanine disaggregation. Together with the observed ligand-induced drLHG4 folding, these results are consistent with the formation of DNA-bound monomeric phthalocyanine. Limited interaction is observed with d(GT)$_7$ that does not fold. The small change in absorption and emission spectra is likely due to some non-selective electrostatic interaction with the unfolded single-strand.

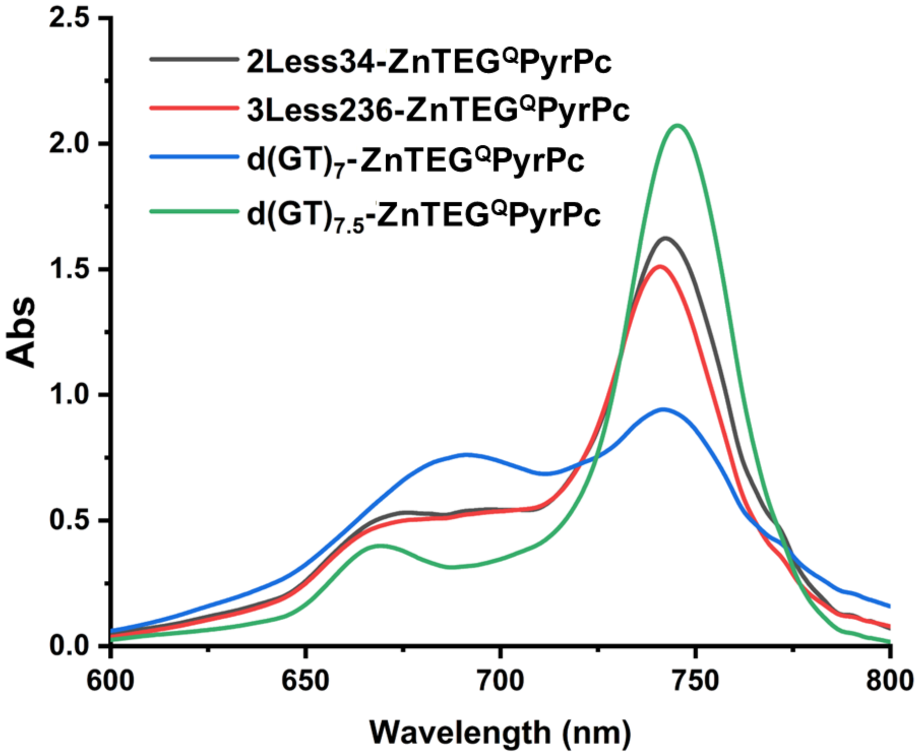


**Figure S21.** UV-visible absorption responses of ZnTEG$^{Q}$PyrPc to selected d(GT)$_{n}$-derived sequences. UV-visible absorption spectra of 5 μM ZnTEG$^{Q}$PyrPc in the presence of 5 μM d(GT)$_{7.5}$, d(GT)$_{7}$, *2Less14* or *3Less236*. drLHG4-competent sequences produced a strong absorption band near 745 nm, consistent with formation of a monomeric ZnTEG$^{Q}$PyrPc-bound state. The response was strongest for d(GT)$_{7.5}$, while d(GT)$_{7}$, which does not form drLHG4 under these conditions, showed a weaker response. These data support preferential productive engagement of ZnTEG$^{Q}$PyrPc with drLHG4-forming sequences.

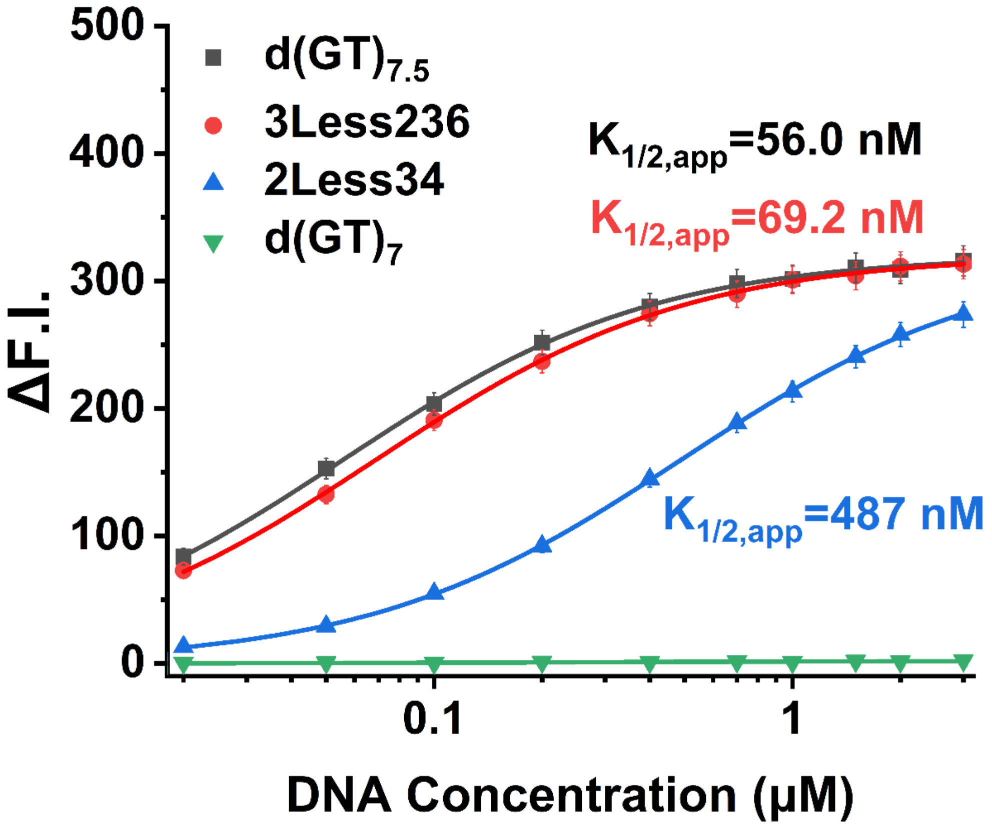


**Figure S22.** Fluorescence titration response of $ZnTEG^{Q}PyrPc$ with selected $d(GT)_n$-derived sequences after mixing and annealing. $d(GT)_{7.5}$ and *3Less236* produced strong fluorescence responses, with apparent half-maximal response concentrations ($K_{1/2,app}$) of 56.0 and 69.2 nM, respectively, whereas *2Less34* produced a weaker response with a $K_{1/2,app}$ of 487 nM. In contrast, $d(GT)_7$ produced little fluorescence response over the same concentration range. Curves represent empirical apparent saturation fits, and error bars indicate SD from three independent measurements. Because $ZnTEG^{Q}PyrPc$ aggregation, monomer stabilization, and DNA binding/folding are coupled in this assay, $K_{1/2,app}$ values should not be interpreted as microscopic 1:1 dissociation constants.

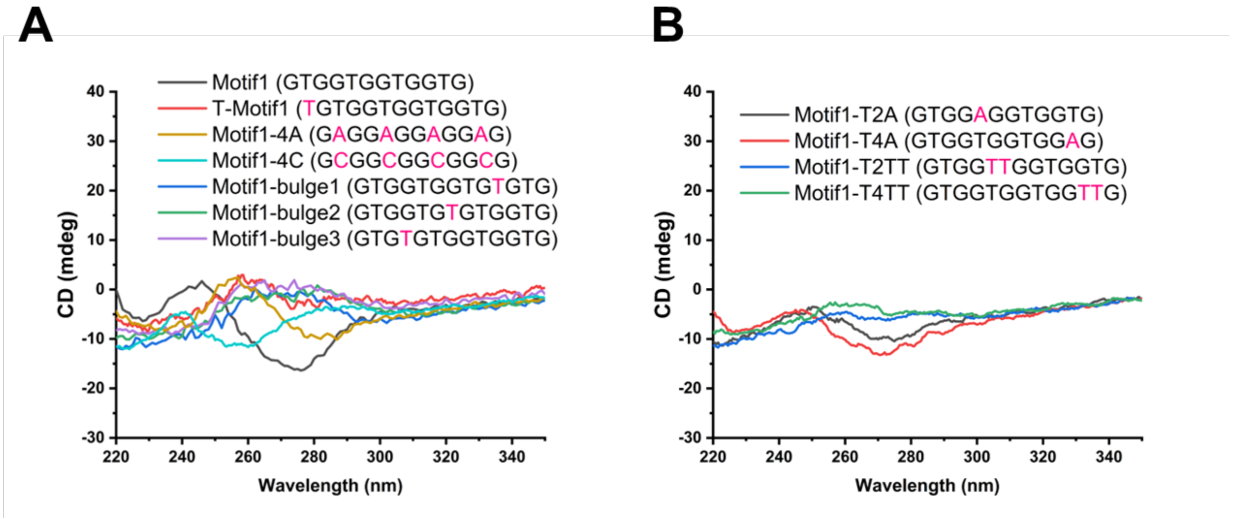


**Figure S23.** Effects of sequence modifications on *Motif1* CD profiles in the absence of dehydrating cosolvents. (A) CD spectra of *Motif1* and variants containing a 5′-terminal T extension (T-*Motif1*), replacement of all four T residues with A or C (*Motif1*-4A and *Motif1*-4C), or a single T insertion at three different positions within G-tracts (*Motif1*-bulge1–3). (B) CD spectra of variants in which the second or fourth T residue of Motif1 was replaced with A (*Motif1*-T2A and *Motif1*-T4A) or expanded to TT (*Motif1*-T2TT and *Motif1*-T4TT). Sequences are shown in the legends, with modified or inserted nucleotides highlighted in magenta.

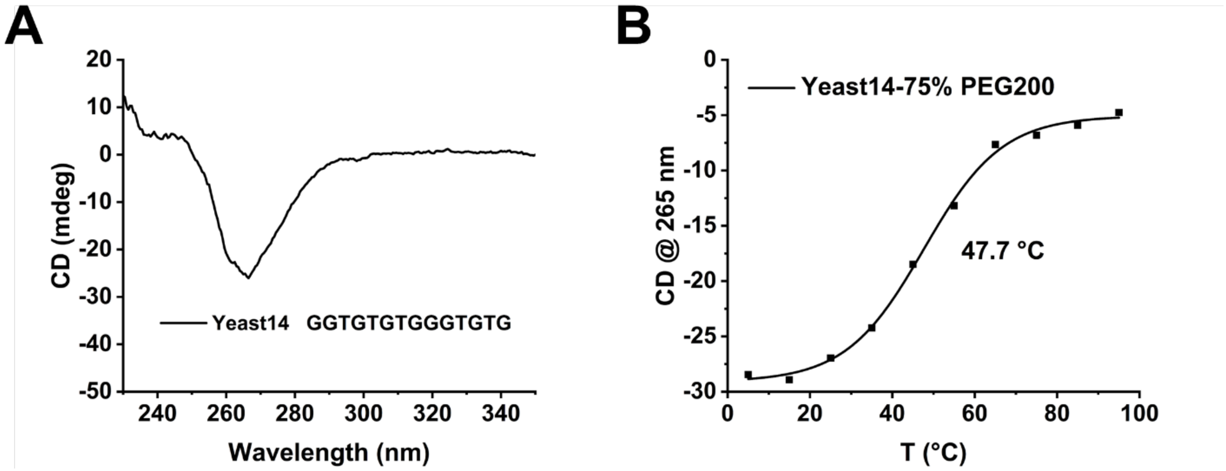


**Figure S24.** Dehydration-induced G4 folding and thermal stability of a budding yeast telomeric-repeat-derived sequence. (A) CD spectrum of Yeast14 (GGTGTGTGGGTGTG) under dehydrating conditions, showing a pronounced negative band near 265 nm. (B) CD melting curve of Yeast14 in 75% (v/v) PEG200, monitored at 265 nm, yielding a $T_m$ of 47.7 °C.

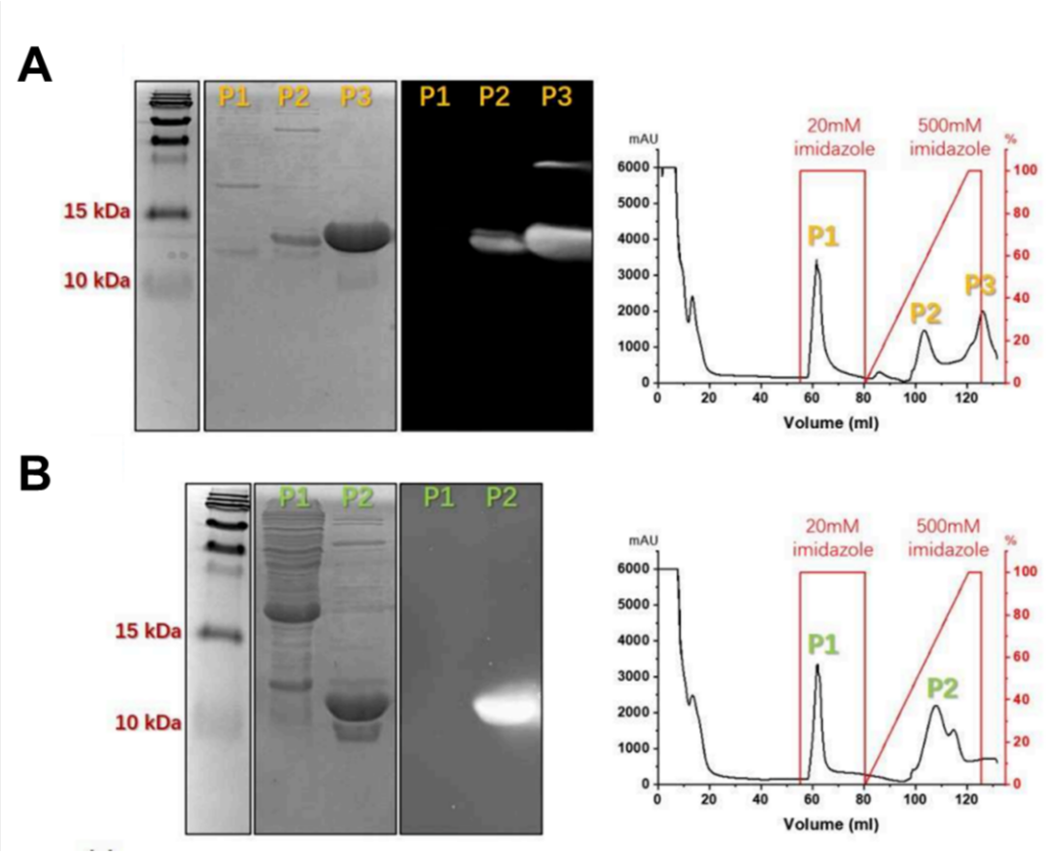


**Figure S25.** Analysis of purification fractions of (A) G4P and (B) NPM-C70. Coomassie Brilliant Blue-stained SDS-PAGE gels (left), Western blots (middle), and purification elution profiles (right) are shown.
